# GLIS3 Marks a Neural-like Progenitor Cell State that Drives Metastasis in Pancreatic Ductal Adenocarcinoma

**DOI:** 10.1101/2025.09.18.677140

**Authors:** Dennis Gong, Jimmy A. Guo, Jennifer Su, Sueda Cetinkaya, Connor Hennessey, Patrick Yu, Ananya Jambhale, Peter L. Wang, Nicholas J. Caldwell, Ashley Lam, Junning Wang, Carina Shiau, Kevin S. Kapner, Laleh Abbassi, Seema Chugh, Shatavisha Dasgupta, Jonathan Nowak, Brian M. Wolpin, M. Lisa Zhang, Mari Mino-Kenudson, Tyler Jacks, Andrew J. Aguirre, William L. Hwang

## Abstract

Pancreatic ductal adenocarcinoma (PDAC) has a high rate of recurrence and metastasis despite intensive therapy. The neural-like progenitor (NRP) transcriptional program is enriched in residual disease after neoadjuvant chemotherapy and radiotherapy; however, the mechanisms for increased NRP expression in the post-treatment setting remain unclear. Here, we find that NRP signatures are strongly enriched in tissue injury and regeneration, and NRP cancer cells co-express transcription factors involved in pancreatic development. We identify both cell autonomous and ligand mediated mechanisms for inducing NRP expression *in vitro*. We establish and characterize isogenic mouse organoid overexpression models for transcription factors associated with the NRP, classical, and basal-like states. We discovered that Glis3 is a key NRP-associated transcription factor that drives malignant properties including greater clonogenicity, tumor growth, and metastasis compared to isogenic models for the classical and basal-like states. Our work highlights the emergence of clinically relevant developmental regeneration programs in the post-treatment context.

## Introduction

Pancreatic ductal adenocarcinoma (PDAC) is an aggressive disease that is frequently detected at advanced stages and responds poorly to conventional treatments^1^. PDAC is dominated genomically by gain-of-function *KRAS* mutations in over 90% of patients^2,3^, and inhibitors of common alleles have entered clinical trials^1^. However, the mutational profiles of PDAC tumors are relatively homogeneous^2,3^, and most mutations do not reliably stratify patients by prognosis or treatment response^4^. Significant attention has therefore been directed to studying transcriptional states in this disease^4–6^.

Gene expression states in pancreatic cancer are associated with distinct prognosis and treatment response^4–9^. Multiple studies have substantiated ‘classical’ (CLS) and ‘basal-like’ (BSL) as the major transcriptional states in untreated pancreatic cancer^4,6,10,11^, with the latter exhibiting more aggressive behavior in patients. Recently, using single-nucleus RNA-seq (snRNA-seq) on primary human tumors, we identified a ‘neural-like progenitor’ (NRP) state in cancer cells that was lowly represented in untreated settings but enriched after neoadjuvant treatment with chemotherapy and radiotherapy (CRT)^7^. Taken together, classical, basal-like, and neural-like progenitor encompass three major cancer cell states in both untreated and previously treated disease.

Despite their associations with clinical outcomes, our functional understanding of these states – particularly the neural-like progenitor subtype – remains insufficient. Neural-like or neuroendocrine gene expression programs have been associated with poor prognosis, treatment resistance, and metastasis in other solid tumors^12,13^. In PDAC, malignant cells exhibit a complex relationship with the nervous system, manifested in part by a high incidence of perineural invasion^14^ and axon guidance pathway alterations^15^. Thus, our identification of the NRP transcriptional program and observations of state enrichment after neoadjuvant treatment motivates investigation of neural gene expression by malignant PDAC cells.

Here, we evaluated the neural-like progenitor state in untreated and treated, primary and metastatic pancreatic cancer and noted its enrichment in metastatic and treatment-resistant disease. We address critical questions regarding the role of reprogramming versus selection in the emergence of NRP and the mechanisms underlying the NRP state in PDAC cancer cells. We provide evidence that NRP can arise from both cell autonomous mechanisms and signals from the post-treatment tumor microenvironment, and that the NRP cell state is poorly modeled *ex-vivo* by conventional patient derived cell lines and organoids. Further, we find that NRP expression is enriched in human and mouse datasets of pancreatitis-induced pancreatic injury and regeneration. We identified *GLIS3* as a key transcription factor (TF) regulator of the NRP state, and developed isogenic mouse organoid models of the classical, basal-like, and neural-like progenitor states by overexpressing state-specific TFs with CRISPR activation. *In vitro* and *in vivo* characterization studies reveal aggressive growth of the basal-like and neural-like progenitor models, as well as a marked metastatic proclivity of the Glis3 driven isogenic model.

## Results

### The neural-like progenitor state is enriched in treated and advanced PDAC

We previously found that neural-like progenitor cancer cells exist at low prevalence (9.9%) in untreated disease but are enriched after neoadjuvant therapy (40.5%) when examining snRNA-seq data derived from primary patient-derived PDAC (**Fig. 1A**). In addition to enrichment of cells annotated as NRP via consensus non-negative matrix factorization (cNMF) scores, the majority of the top 200 weighted cNMF NRP genes were also enriched in residual disease (**Fig. 1A**).

**Figure 1:**
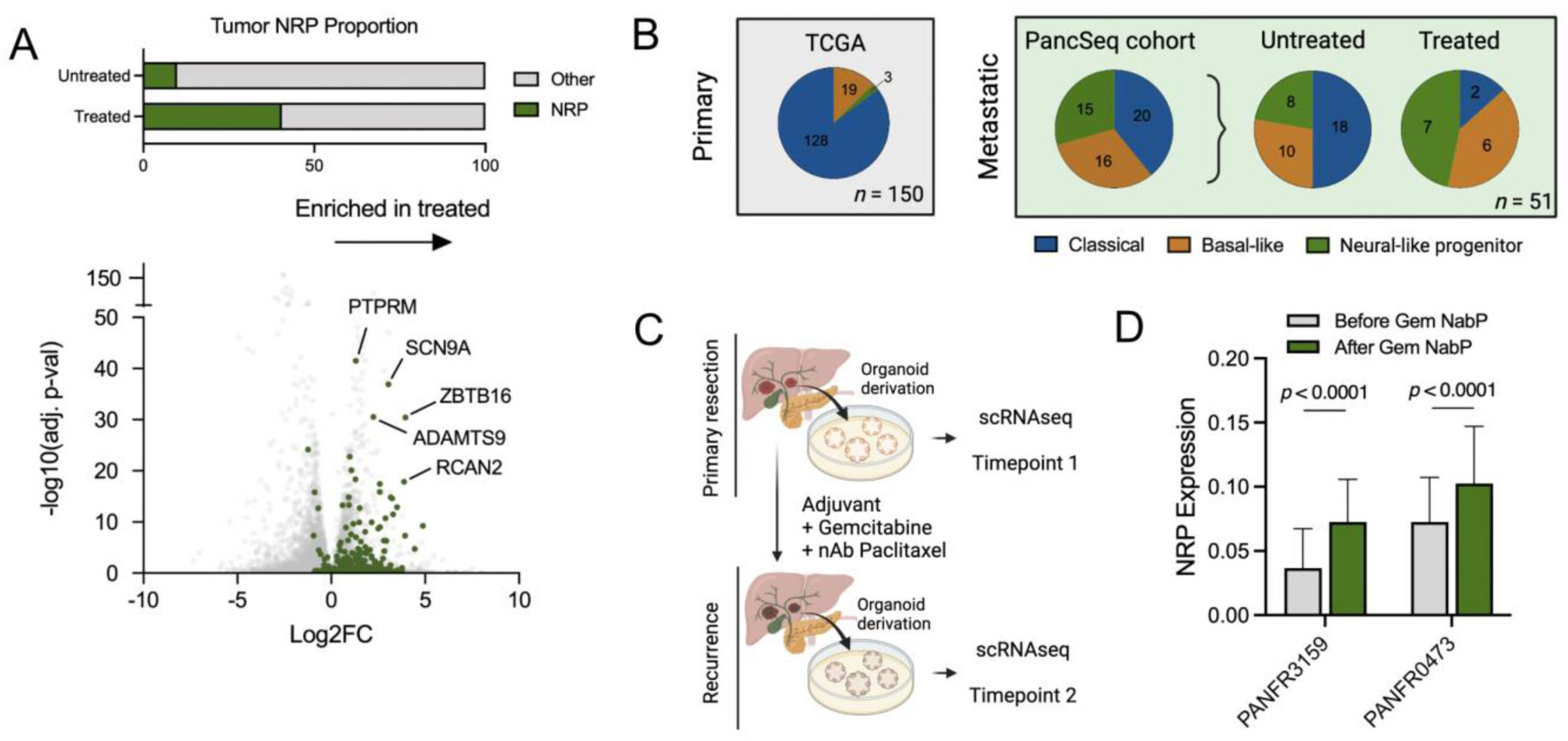
The neural-like progenitor state is enriched in post-CRT treated pancreas cancer specimens. (a) Schematic of NRP cell enrichment in residual treated tumor (top) and scatterplot of differentially expressed genes comparing treated versus untreated patients (bottom) in our snRNAseq cohort^7^. Green colored dots represent genes in the top 200 cNMF weighted genes in the NRP signature. NRP cells expanded as a proportion of cancer cells from 9.9% to 40.5%. (b) Pie chart representing proportion of patients in the TCGA cohort of primary PDAC with dominant expression of each subtype (left) and pie charts representing proportion of patients in PancSeq metastatic PDAC cohorts with dominant expression of each subtype (right). (c) Study schematic for organoid derivation and scRNAseq experiment using patient matched pre- and post-treatment organoids. (d) Grouped barplot depicting scRNAseq profiling of NRP signature expression in organoids derived from patients before and after treatment with gemcitabine and nab-paclitaxel (Gem NabP) multi-agent chemotherapy.

However, the relevance of the NRP state in other patient cohorts and beyond the localized setting remains incompletely defined. To answer this question, we assembled three independent single-cell RNA-seq (scRNA-seq) datasets^16–18^ encompassing 95 unmatched primary PDAC patients undergoing surgical resection (47 treated with neoadjuvant therapy, 48 untreated). We scored annotated epithelial cancer cells for NRP^7^, BSL^6^, and CLS^6^ gene programs and performed a sample size bootstrapped Mann-Whitney test to compare subtype signature expression between treated and untreated samples. In aggregate, we identified higher NRP scores in samples treated with neoadjuvant therapy, with variable impact on CLS or BSL expression (**Fig. S1A**).

In analyzing bulk RNAseq data from the untreated TCGA PDAC cohort^3^, we noted that the NRP state was the predominant malignant subtype in fewer than 3% of primary pancreatic tumors; most were classified as CLS or BSL (**Fig. 1B**). However, we found increased NRP expression in persister cells after neoadjuvant therapy in a separate cohort of 97 bulk RNA sequenced samples^19^ (**Fig. S1B,C**), consistent with our prior results^7^. Interestingly, we found higher expression of NRP in patients treated with FOLFIRINOX vs gemcitabine backbone regimens (*p* = 0.0015; Mann–Whitney U test) (**Fig. S1B,C**), with both treated categories having numerically increased expression of NRP relative to untreated samples, and statistically significant enrichment in the FOLFIRINOX treated group (*p* < 0.0001; Mann–Whitney U test).

To investigate the prevalence of the NRP state in disseminated disease, we classified a subset of metastatic biopsies (n=51) from the PancSeq^20^ patient cohort as predominantly CLS, BSL, or NRP. In contrast to its low relative representation in the primary TCGA cohort, approximately one-third of PancSeq biopsies were NRP predominant relative to the classical and basal-like states (**Fig. 1B**). We further asked whether chemotherapy also alters the prevalence of states in the metastatic setting. Indeed, treated PancSeq biopsies had significantly higher representation of NRP and BSL, and lower representation of CLS compared to untreated biopsies (**Fig. 1B**). Taken together, these data suggest that the NRP state is also enriched after cytotoxic treatment in more advanced disease.

Two of the PancSeq biopsies were patient-matched pairs pre- and post-treatment with gemcitabine plus nab-paclitaxel, which provided a unique opportunity to directly examine for treatment-induced state changes. To this end, we established matched pre- and post-treatment patient-derived organoids (PDOs) from these biopsies and performed scRNAseq to measure gene expression differences (**Fig. 1C**). We observed significantly higher NRP expression in post-treatment organoids (*p* < 0.0001; Mann–Whitney U test) (**Fig. 1D**).

To assess for potential co-expression of distinct states within individual cells, we performed correlation analyses across all epithelial cancer cells in our snRNA-seq and scRNA-seq datasets^7^. We observed correlation across various basal and mesenchymal gene sets, as well as correlation across gene sets for classical or glandular PDAC cells (**Fig. S2A**). NRP expression had low or inverse correlation with all other gene sets except a PDAC neuroendocrine signature^7^. We also compared the pancreatic NRP signature^7^ to neural and neuroendocrine transdifferentiation states described in other malignancies, including small cell lung cancer and prostate cancer^21,22^ (**Fig. S2B,C**). Gene set overlap and single-cell correlation analyses revealed modest concordance among NRP and neuroendocrine-like states in PDAC and small cell lung cancer.

In summary, we have verified the presence of the neural-like progenitor gene expression program in primary and metastatic PDAC with enrichment after cytotoxic treatment. Furthermore, we found that the NRP transcriptional signature shares certain genes associated with previously reported neural and neuroendocrine signatures in other malignancies.

### Pancreatic injury is associated with neural-like progenitor expression programs

Functional study of transcriptional states necessitates representative and tractable model systems. However, culture-based conditions are known to bias towards specific cell states^6,8^. For instance, PDAC cell lines tend to be over-represented with basal-like characteristics, while organoids tend to be enriched in classical features^6,8^. The representation and stability of the NRP state in existing *in vitro* and *ex vivo* models is unknown. To this end, we scored human pancreatic cancer cell lines and PDOs with accompanying RNA-seq data (**Fig. S3A**) for the CLS, BSL, and NRP signatures and compared them to their representation in patient-derived tissue specimens. Most PDAC models did not express the NRP state at high levels with a few exceptions (e.g., PANFR0202 T2, PACADD165, PACADD137) (**Fig. S3A**). The model most transcriptionally similar to the tissue-derived NRP state was a PDO derived from a post-treatment PDAC metastatic biopsy (PANFR0202 T2).

To evaluate the stability of this PDO in culture over time, we analyzed scRNAseq data of PDAC organoids at various passage points. NRP expression was variable across passage points and in some instances strongly decreased with subsequent passages (**Fig. S3B**), suggesting that standard organoid culture conditions may be insufficient for maintaining the NRP state over time. These data indicate that the NRP state is not well represented or maintained in current PDAC cell line and organoid models, which may contribute to our limited understanding of NRP and neuroendocrine states in cancer more broadly. Indeed, the lack of persistence in culture of neuroendocrine state models in prostate cancer has been similarly described^23^.

In our previous work, we had observed that NRP expression correlates with treatment response^7^, with the strongest NRP expression in tumors with the greatest response to therapy and therefore the least amount of residual disease. Given that the NRP state is enriched post-treatment (**Fig. 1A-D**), we sought to determine whether the NRP state could be rescued in culture by *ex vivo* treatment. Surprisingly, we found that chemotherapy did not significantly affect the expression of most NRP signature genes profiled immediately after treatment *in vitro*, both in 2D mouse *Kras*^LSL-G12D/+^;*Trp53*^FL/FL^ (KP) cell lines and in several established PDOs (**Fig. S4A,B**). Given the lack of NRP program expression in our reductionist *in vitro* systems, we proceeded to investigate whether more sophisticated *in vivo* model systems are required to observe signature expression.

The 200 gene NRP signature^7^ contains not only genes related to neurogenesis (*p* = 5.01e-16) and neuronal development (*p* = 1.00e-16) but is also highly enriched for fetal (*p* = 2.50e-18) and physiologic pancreatic ductal signatures (*p* = 1.68e-42) (**Fig. S5A**). While several of the NRP signature genes are also highly expressed in non-transformed ductal epithelial cells (**Fig. S5B**), the expression of these genes in cancer cells distinguishes them from other cancer cells. However, the shared expression of some genes with physiologic epithelium suggested that NRP may represent a common regenerative program expressed by both injured healthy and malignant epithelia. We hypothesized that chemotherapy and radiotherapy may be acting similarly to other sources of tissue damage, inducing expression of a conserved regenerative epithelial cell state.

To test this hypothesis, we first turned to pancreatic injury models and scored NRP expression in regenerating epithelial cells. Indeed, we identified increased NRP expression (NES: 1.998, *p* < 0.0001) in epithelial spheroids derived from mice with caerulein-induced pancreatitis^24^ (**Fig. 2A**) but NRP expression decreased markedly with prolonged culture similar to prior findings with PDOs. In a separate dataset^25^, we also observed a temporal increase in NRP expression by ductal cells *in vivo* over the course of caerulein-induced pancreatitis (**Fig. 2B**). NRP expression was slightly lower at 14 hours after induction of pancreatitis, but subsequently rebounded, peaked at 5 days, and remained elevated relative to baseline at 14 days (Sen’s slope *p* = 3.7e-10).

**Figure 2:**
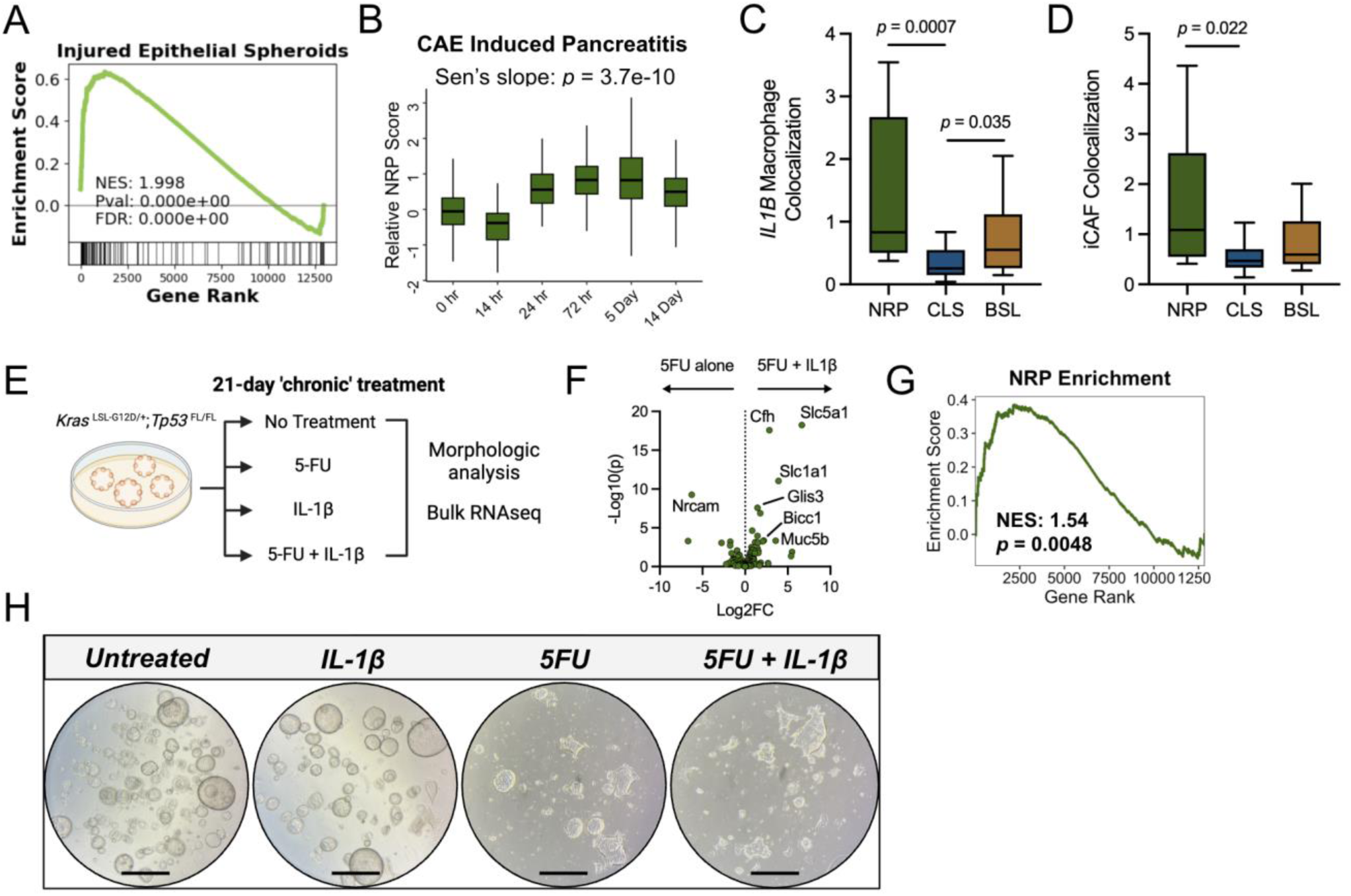
Neural-like progenitor expression programs are observed specifically in pancreatitis and injury repair. (c) GSEA enrichment plot of NRP signature in epithelial spheroids derived from a CAE induced pancreatitis model compared to uninjured baseline^24^. (d) Boxplot depicting NRP score in CAE induced pancreatitis mouse model assayed via scRNAseq at acute timepoints^25^. (c,d) Quantification of colocalization of NRP cancer cells with *IL1B* expressing macrophages (c) and (d) inflammatory CAFs^66^. (e) Schematic of chronic inflammation treatment model to assay effects of recombinant IL-1β and 5-FU on murine PDAC organoids. (f) Volcano plot depicting change in NRP gene expression in 5-FU treated organoids with IL-1β treatment versus 5-FU alone. (g) GSEA plot showing enrichment of NRP signature genes in RNAseq of 5-FU and IL-1β treated organoids versus 5-FU treated organoids. (h) 4x Phase imaging of organoids following treatment. Scale bars represent 1,000 µm.

These experiments were performed in wild-type mice, but most pancreatic cancers arise in the context of gain of function KRAS mutations. Thus, we queried NRP expression in a published scRNAseq dataset^26^ of caerulein-induced pancreatitis and PanIN development in mice harboring pancreas specific KRAS^G12D^. We identified markedly higher NRP state expression in pancreatic ductal cells after prolonged pancreatitis at 3 weeks post caerulein, relative to immediately after induction at 1 or 2 days post treatment (*p* < 2.2e-16; Mann–Whitney U test) (**Fig. S5C**). Additionally, we observed that NRP expression increased throughout the development and progression of pancreatic intraepithelial neoplasia (PanIN) across a 27-week period with exposure to pancreas specific KRAS^G12D^, relative to healthy non-mutant baseline (**Fig. S5D**).

Our findings in mouse models were supportive of our hypothesis that NRP is enriched in epithelial cells subjected to tissue damage. Next, we asked if we could also identify NRP expression in the injury or stress context within human tissues. We noted that several NRP genes such as *SPP1, CRP,* and *MUC5B* overlapped with characteristic markers of PanIN-like atypical ductal cells^7,16^ (**Fig. S5B**). Additionally, NRP expression is higher in non-malignant adjacent pancreas tissue (TCGA) compared to physiologic pancreas tissue (GTEx) (*p* = 1.66e-13; Mann–Whitney U test) (**Fig. S5E**), indicating that non-malignant tissue in the peritumoral environment can upregulate injury-associated NRP signatures. Furthermore, we identified striking overlap between the NRP signature and marker genes of a ‘stressed progenitor-like’ pancreatic ductal cell^27^ (**Fig. S5F**) (*p* =1.80e-20; Fisher’s Exact test) including genes associated with canonical ductal lineage (e.g., *CFTR, SLC4A4*), tissue repair (e.g., *SPP1*), inflammatory response (e.g., *CRP*), cell proliferation and migration (e.g., *PDGFD*), angiogenesis (e.g., *HIF1A*), and development (e.g., *ONECUT2*, *MEIS2*). NRP cancer cells strongly express the pancreatic progenitor marker *DCLK1*, which was recently reported to be highly expressed in a poorly differentiated population of PDAC cells exhibiting neuronal lineage priming^28^ (*p* < 0.0001; Mann– Whitney U test) (**Fig. S5G**).

Altogether, our analyses nominate NRP as a tissue repair program conserved across malignant and physiologic ductal cells. Additionally, we provide evidence that microenvironmental stress (e.g., pancreatitis, adjacent cancer, drug treatment) can induce expression of the NRP state.

### The inflammatory tumor microenvironment promotes the neural-like progenitor state

We next sought to understand the mechanisms by which cellular stress is capable of inducing NRP gene expression. Given the lack of NRP induction in the simplified *in vitro* setting (**Fig. S4A,B**), we hypothesized that NRP expression may require microenvironmental factors such as soluble ligands to be expressed. Returning to our *in vitro* chemotherapy treated cells, we noted that proinflammatory ligands capable of recruiting immune cells (e.g., *Lcn2*, *Csf1, Cxcl5, Ptgs2*) were among the most upregulated upon short-term treatment with 5-FU based chemotherapy (**Fig. S4A,B**), consistent with previous reports^29^. DNA damaging agents induce cellular stress responses through several mechanisms including DNA damage induced senescence and release of senescence associated secretory factors (SASP), release of damage associated molecular patterns (DAMPs), and signaling through NF-kB, P38-MAPK, and related pathways^30^. Chemotherapy has previously been reported to increase macrophage infiltration *in vivo* by inducing *Csf1* expression in malignant cells^31^ and interestingly, NRP cells express higher levels of *CSF1* than other transcriptional subtypes in our snRNA-seq data^7^.

To investigate the gene expression profiles of cell types commonly present in the microenvironment of NRP cells, we leveraged our recently published cohort of 13 primary PDAC tumors profiled with 990-plex spatial molecular imaging^32^. We utilized our previous annotation schema classifying two subtypes of cancer associated fibroblasts (e.g., myCAF, iCAF) and three subtypes of malignant cells (e.g., CLS, BSL, NRP). Additionally, we annotated *IL1B* expressing macrophages given recent reports describing *IL1B*^+^ macrophages as potent mediators of pathogenic inflammation^33^. In analyzing our spatial transcriptomics data, NRP glands were smaller than classical glands and similar in cluster size to basal-like cells that typically form small aggregates or sheets (**Fig. S6A**). NRP cells were enriched in communities with iCAFs and *IL1B*+ macrophages (**Fig. 2C,D**), consistent with our hypothesis that the NRP state is supported by an inflammatory microenvironment. Strikingly, we observed tissue repair programs in CAFs and macrophages as most differentially expressed in cells near versus far away from NRP cells (**Fig. S6B,C**). These included MHC complex (*p* = 2.93e-14) and antigen presentation (*p* = 3.39e-10) gene sets in CAFs, matrix remodeling (*p* = 9.45e-7) gene sets in macrophages, and pro-regenerative *IL-4*/*IL-13* signaling in both CAFs (*p* = 6.24e-5) and macrophages (*p* = 1.40e-7). Transcriptionally, we found that NRP and iCAF cells have enriched expression of macrophage recruitment and persistence factors *CCL2* and *CSF1* relative to other subtypes, and *IL1B* expressing macrophages had highest expression of receptors *CCR2* and *CSF1R* relative to other macrophages, presenting the possibility that NRP and iCAF cells recruit tumor associated macrophages in a feedforward loop^33^.

Finally, we sought to determine whether we could induce the expression of NRP genes in PDAC organoids. We hypothesized that specific ligands secreted by non-malignant microenvironmental cells could induce expression of NRP genes in cancer cells. IL-1β has been shown to induce iCAF polarization^34^, promote chemoresistance through NF-kB signaling^35^, increase organoid formation proficiency *in vitro*, and stimulate tumor growth *in vivo*^33^. Interestingly, NRP cells also have the highest expression of *IL1R1*, which encodes the cognate receptor for IL-1β. Thus, we treated mouse KP organoids with recombinant IL-1β over the course of 3 weeks with or without 5-FU to evaluate cell state changes both transcriptionally with RNA-seq and morphologically with phase contrast imaging (**Fig. 2E**). Consistent with our hypothesis, treatment of mouse KP organoids using IL-1β with or without 5-FU led to enrichment of NRP signature genes including the neuroendocrine developmental transcription factor *Glis3*, solute transporters *Slc5a1* and *Slc1a1*, and the RNA binding protein *Bicc1* which was previously associated with PDAC stemness and chemoresistance^36^ (**Fig. 2F,G**). While organoids treated with IL-1β alone generally maintained the same luminal morphology as unperturbed KP organoids, the morphology changed with the addition of 5-FU. Chemotherapy treated conditions had smaller organoids and some morphologic heterogeneity but changed most markedly with the combination of 5-FU and IL-1β, characterized by greater eccentricity and heterogeneity (**Fig. 2H**).

We previously observed that a PDO line treated *ex vivo* with chemotherapy and radiation (PDAC_U_12) followed by a three-day rest period exhibited enriched NRP expression without exogenous cytokine supplementation^7^, suggesting additional involvement of cell-autonomous mechanisms. This finding is in contrast to the lack of NRP enrichment immediately after treatment of human and mouse organoids with chemotherapy (**Fig. S4A,B**). Indeed, when we directly compared mouse KP organoids rested three days after 5-FU treatment to those analyzed immediately after treatment, we observed significantly higher NRP expression in the rested organoids (**Fig. S4C-E**). Moreover, rested organoids were less proliferative and expressed fewer genes involved in drug induced DNA repair but had enriched expression of gene sets related to neural functions, inflammatory response, membrane lipid metabolism, hypoxia, and cell adhesion (**Fig. S4F**). Thus, rest and recovery after cytotoxic injury is another critical factor for inducing NRP expression, consistent with our observations in pancreatitis (**Fig. S5C,D**).

In summary, our data suggests that NRP represents a coordinated and conserved endogenous pancreas tissue repair program that integrates both cell autonomous and non-autonomous signals from inflammatory microenvironmental cells such as *IL1B*+ macrophages and inflammatory CAFs. Moreover, our results suggest that NRP state expression may reflect stress-induced epithelial reprogramming, as well as development of tumor cell-intrinsic features that drive chemoresistance.

### Inferring transcription factors associated with the neural-like progenitor state

Detailed characterization of the NRP state in comparison to other malignant states in PDAC requires high fidelity, stable, and experimentally tractable model systems. Given the challenges of maintaining state-specific gene expression profiles in standard culture-based conditions (**Fig. S3A,B**), we reasoned that we could stabilize and therefore study these states in a systematic manner by generating isogenic models through TF overexpression. Additionally, the top 200 NRP cNMF genes were significantly enriched for development associated TFs (**Fig. 3A**)^7^. While key TF regulators of the classical (*GATA6*) and basal-like (*deltaNTrp63*) states have been described^37,38^, those of the NRP state are unknown, precluding the generation of isogenic NRP model systems. As such, we performed *in silico* analyses to infer candidate TF regulators of the NRP state.

**Figure 3:**
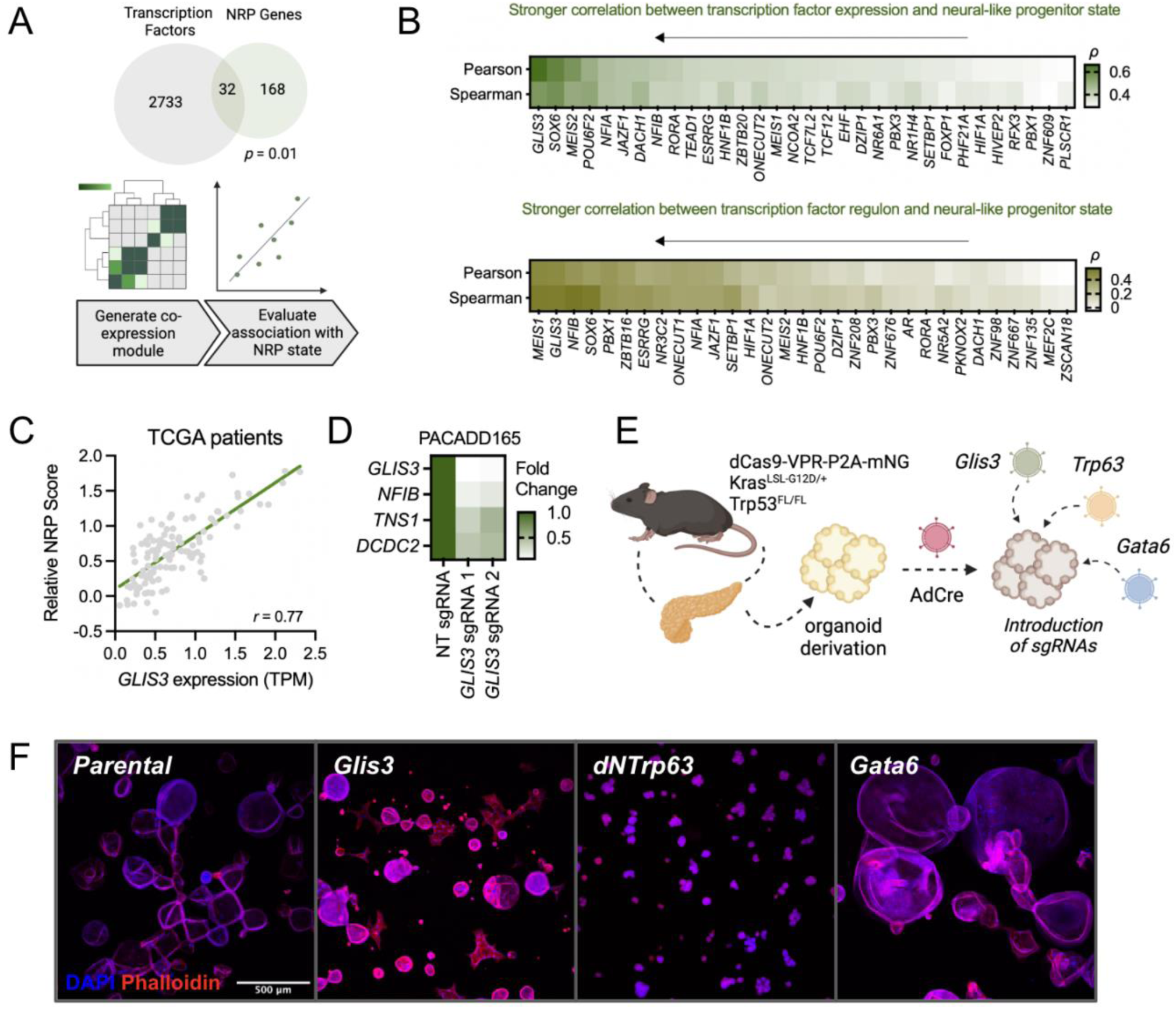
Identification of TF regulators of the neural-like progenitor state. (a) Overlap of transcription factors and top 200 NRP genes by cNMF weights^7^. Statistical significance of overlap calculated by Fisher’s exact test. (b) Heatmaps of Pearson and Spearman correlation coefficients between single gene transcription factor RNA expression (top) and signature score of the NRP state, and transcription factor regulon (ie. target genes of each transcription factor) expression and NRP signature (bottom) across malignant cells from a single-nucleus RNA-seq dataset of human primary PDAC. (c) Correlation of GLIS3 expression with NRP signature expression in TCGA bulk RNAseq data. (d) qPCR of select NRP genes after knockout of GLIS3 from patient-derived organoid model. (e) Schematic of pancreatic organoid derivation from Kras^LSL-G12D/+^; p53^FL/FL^ altered mice, Cre recombination, and CRISPRa cassette integration. Isogenic CRISPRa organoids (Kras^LSL-G12D/+^;Trp53^FL/FL^;R26-dCas9-VPR-mNG) were engineered with gRNAs targeting promoter regions of Gata6, dNTrp63, and Glis3. (f) Confocal immunofluorescence imaging of isogenic organoid lines grown in Matrigel matrix (**Methods**). Scale bar of 500 µm is the same for all panels.

First, we reasoned that the abundance of a TF regulator should correlate with state expression (**Fig. 3B**). To this end, we computed correlation coefficients between candidate TFs within the top 200 cNMF-weighted genes and NRP expression in malignant cells from the snRNA-seq cohort^7^. Among these candidates, *GLIS3* ranked as the most correlated TF (**Fig. 3B *top***). Other correlated TFs included several known to interact with each other, such as *NFIB* and *NFIA*, as well as *PBX1*, *MEIS1*, and *MEIS2*. *NFIB,* in particular, has been identified as the driver of a pro-metastatic neuronal expression signature in small cell lung cancer^39^.

Next, we reasoned that the regulon activity of *bona fide* NRP TFs should also correlate with state expression. We therefore leveraged a previously described method called SCENIC^40^ to infer regulons for each candidate TF using our snRNA-seq data^7^, and computed correlation coefficients between expression of the NRP state and TF regulons in malignant cells (**Fig. 3B *bottom***). We observed concordance with our prior TF expression analysis, as *MEIS1*, *GLIS3*, and *NFIB* were the three most highly-ranked candidates by this regulon-based approach (**Fig. 3B**). We identified *GLIS3* as a top candidate in both analyses, and reassuringly we found that *GLIS3* expression also correlated most strongly with overall NRP signature expression in bulk RNA sequencing data from the PDAC TCGA cohort (**Fig. 3C**), and that *Glis3* expression was induced upon IL-1β treatment in our prior organoid experiments (**Fig. 2H**). To determine the necessity of *Glis3* for maintaining key cell state genes, we performed a CRISPR knockout experiment by transducing a *Glis3* targeting guide RNA into PACADD165, a Cas9 expressing patient-derived organoid with high NRP expression, which reduced the expression of cell state markers *Glis3*, *Nfib*, *Tns1*, and *Dcdc2a* (**Fig. 3D**).

We then sought to validate *Glis3* as a regulator of the NRP state, and generated independent isogenic mouse models of the CLS, BSL, and NRP states. We leveraged a previously described *Kras*^LSL-G12D/+^;*Trp53*^FL/FL^ organoid model^41^ encoding a germline dCas9-VPR CRISPR activation (CRISPRa) system for facile and modular overexpression of transcription factors at their endogenous regulatory loci (**Fig. 3E**). We transduced parental organoids with lentivirus containing sgRNAs targeting the promoter regions of *Gata6*, *dNTrp63*, and *Glis3* to generate isogenic models of the CLS, BSL, and NRP states, respectively. We chose *Gata6* and *dNTrp63* due to their previously described functional associations with the classical and basal-like cell states,^38,42^respectively.

We measured the transcript and protein levels of each respective target gene by quantitative PCR (qPCR) and Western blot, respectively, to confirm endogenous upregulation relative to the parental line (**Fig. S7A**). Indeed, we observed pronounced upregulation of *Gata6*, *dNTrp63*, and *Glis3* transcript expression in their respective models. Downstream of TF upregulation, we noted appropriate expression changes in state-specific marker genes: *Dcdc2a* was upregulated 38-fold in the *Glis3* model, *Tff1* 28-fold in the *Gata6* model, and *Krt17* 1897-fold in the *dNTrp63* model (**Fig. S7A**), suggesting that genetic manipulation of cell-intrinsic factors can indeed induce state expression. Given that culture-based conditions can lead to NRP state shifts over time (**Fig. S3B**), we sought to evaluate whether cell-intrinsic factors could maintain states after extended passage. As such, we performed qPCR of NRP marker genes 10 passages after the initial time point at which they were profiled (**Fig. S7B**). Respective cell state markers remained activated at the later time point (**Fig. S7B**), indicating the relative stability of these genetically engineered models in culture.

To perform a global assessment of transcriptional changes induced by each TF, we performed RNA-seq on these isogenic CRISPRa organoids. As expected, we observed that the *Gata6* model most closely represented the classical state, the *dNTrp63* model the basal-like state, and the *Glis3* model the NRP state (**Fig. S7C**). Activated genes specific to the NRP model included *Glis3*; *Pcp4l1*, a Purkinje cell gene expressed in the central nervous system^43^; *Nat8l*, a brain-specific gene that catalyzes the synthesis of N-acetylaspartate^44,45^; *Dclk1*, a doublecortin-like kinase gene that demarcates tuft cells^46^; *Spp1*, a gene linked to tumor invasiveness^47^; and *Shank2*, a postsynaptic density scaffolding gene^48^. Relative to the parental organoid, the Glis3 organoid line was enriched in gene sets related to cell motility and migration with purportedly greater responsiveness to exogenous stimuli such as chemokines and oxygen gradients (**Fig. S7D**).

Activated genes specific to the classical model included *Tff3*, a trefoil factor gene involved in the maintenance of gastrointestinal mucosa^49^; and *Muc5ac*, a gel-forming mucin gene^50^. Genes specific to the basal-like model included those reminiscent of epidermis development or squamous morphology, such as *Krt5*, *Krt6a*, *Krt17, Col17a1*, and *Dsg3*, as well as *Tgfbi*, a TGF-beta induced gene. Gene sets enriched in the *Gata6* model related to glutathione metabolism, and gene sets enriched in the *dNTrp63* model related to keratinization and epidermal differentiation (**Fig. S7E,F**).

Thus, we have identified *Glis3* as a transcription factor strongly co-expressed with the NRP signature in both *in silico* and functional experiments. Isogenic organoids engineered with CRISPRa based transcription factor activation enabled sustained expression of cell state associated genes and transcriptional signatures.

### Glis3 overexpression increases clonogenic, migratory, and invasive capacity

To explore the phenotypic features of our isogenic subtype-specific organoids, we first performed confocal immunofluorescence and phase contrast imaging (**Fig. 3F, S8A**). Morphologically, the Gata6-driven organoids resembled their parental counterparts; the dNTrp63-driven organoids exhibited luminal filling reminiscent of squamous and basal-like tissue morphology; and the Glis3-driven organoids featured spindle-like projections, similar to our findings in 5-FU and IL-1β treated KP organoids (**Fig. 2H**). We also noted in human organoids that transcriptionally resemble classical or basal-like identities, classical organoids share the spherical luminal shape of the Gata6-driven model, while the basal-like human organoids share morphologic resemblance with both the Glis3 and dNTrp63-driven models with spindle-like projections and luminal filling (**Fig. S8B**). Consistent with our observations in human PDAC^32^ tissue, the Glis3-driven NRP and dNTrp63-driven basal-like models grew smaller but more abundant organoid colonies (*p* < 0.001; Mann–Whitney U test) compared to the parental and Gata6-driven classical lines (**Fig. S6A, 4A,B**). Moreover, we observed the highest clonogenicity in the Glis3 model, compared to the dNTrp63 model (*p* = 0.0022; Mann–Whitney U test) and Gata6 model (*p* = 0.0022; Mann–Whitney U test) in both 2D and 3D formats (**Fig. 4A,C**), consistent with the Glis3-driven NRP state exhibiting a progenitor-like phenotype.

**Figure 4:**
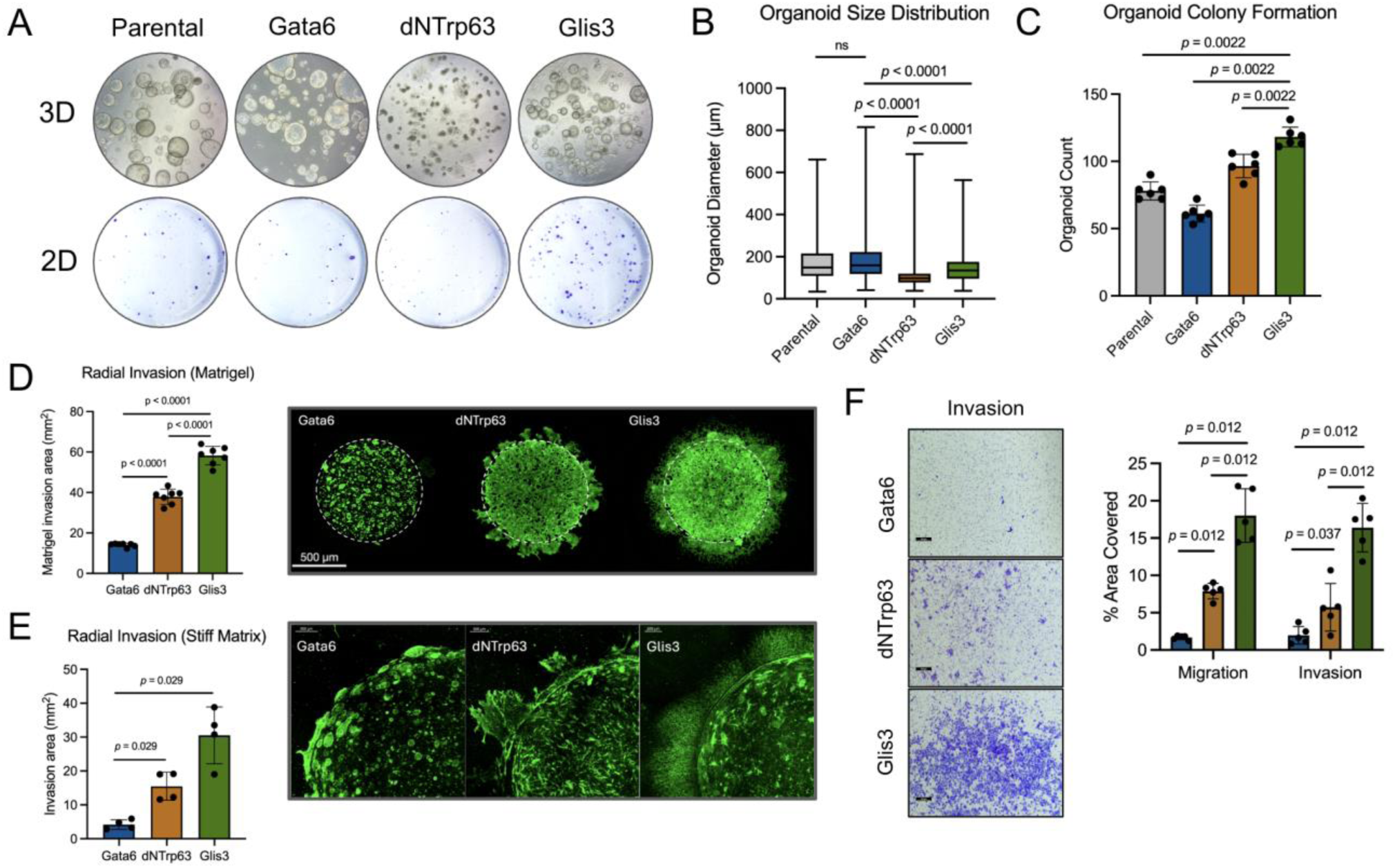
The neural-like progenitor state exhibits stem-like properties and is inducible in vitro. (a). Representative images of colony formation assays in 3D (top) and 2D (bottom) using isogenic cell lines with quantification of (b) size and (c) colony count from 3D assays. (d). Radial invasion assay and quantification comparing isogenic lines in soft Matrigel assay format and in (d) stiff collagen I matrix. Invasion area was quantified as cells invading out of seeded plug. Representative images of invading cells are shown on the right. (f) Boyden chamber migration/invasion assay with associated quantification. Representative fields of view for each line are shown.

Next, we assessed the migratory and invasive properties of our isogenic models. We designed a 3D radial invasion assay wherein cells are centrally seeded in a lower density Matrigel and invade radially through a higher density Matrigel (**Fig. 4D**) or collagen I matrix (**Fig. 4E; Methods**). Glis3 driven lines invaded through both the Matrigel (*p* < 0.0001; Mann–Whitney U test) and collagen I matrix (*p* < 0.01; Mann–Whitney U test) at a faster rate than both the Gata6 and dNTrp63 models. Consistent with the more aggressive behavior of basal-like PDAC in humans compared to the classical subtype^51,52^, the dNTrp63-driven line had significantly higher invasive capacity than the Gata6-driven organoid line (**Fig. 4D,E**). These cell state-associated differences in migration and invasion were recapitulated with the 2D Boyden chamber assay using isogenic TF-driven cell lines instead of organoids (**Fig. 4F**).

Thus, we have observed by several orthogonal approaches that Glis3-driven NRP cell lines and organoids exhibit greater clonogenic, migratory, and invasive capacity than their Gata6-driven CLS and dNTrp63-driven BSL counterparts.

### Glis3 promotes tumor growth and metastasis *in vivo*

To study these malignant cell states *in vivo*, we orthotopically transplanted our isogenic CRISPRa organoid lines into syngeneic host mice (**Fig. 5A; Methods**). To evaluate the tumor growth kinetics of each organoid line, we harvested and weighed primary tumors at 5 weeks, 8 weeks, and 11 weeks and surveyed for liver and lung metastases 11 weeks after transplantation (**Fig. 5A**). Histopathologic examination of tissue sections by three board-certified GI pathologists (N.J.C., J.N., M.L.Z., & M.M-K.) confirmed *bona fide* formation of adenocarcinomas and revealed that the morphology of mouse *Glis3*-, *Gata6*-, and *dNTrp63*-activated tumors were reminiscent of human neural-like progenitor, classical, and basal-like tumors, respectively (**Fig. 5B**). In particular, *dNTrp63*-activated tumors had greater area with poorly differentiated histology, whereas *Gata6-*activated and parental tumors featured more moderately to well-differentiated areas (**Fig. 5C**). The *Glis3*-activated tumors had roughly equal proportion of poorly and moderately differentiated tumor areas (**Fig. 5C**).

**Figure 5:**
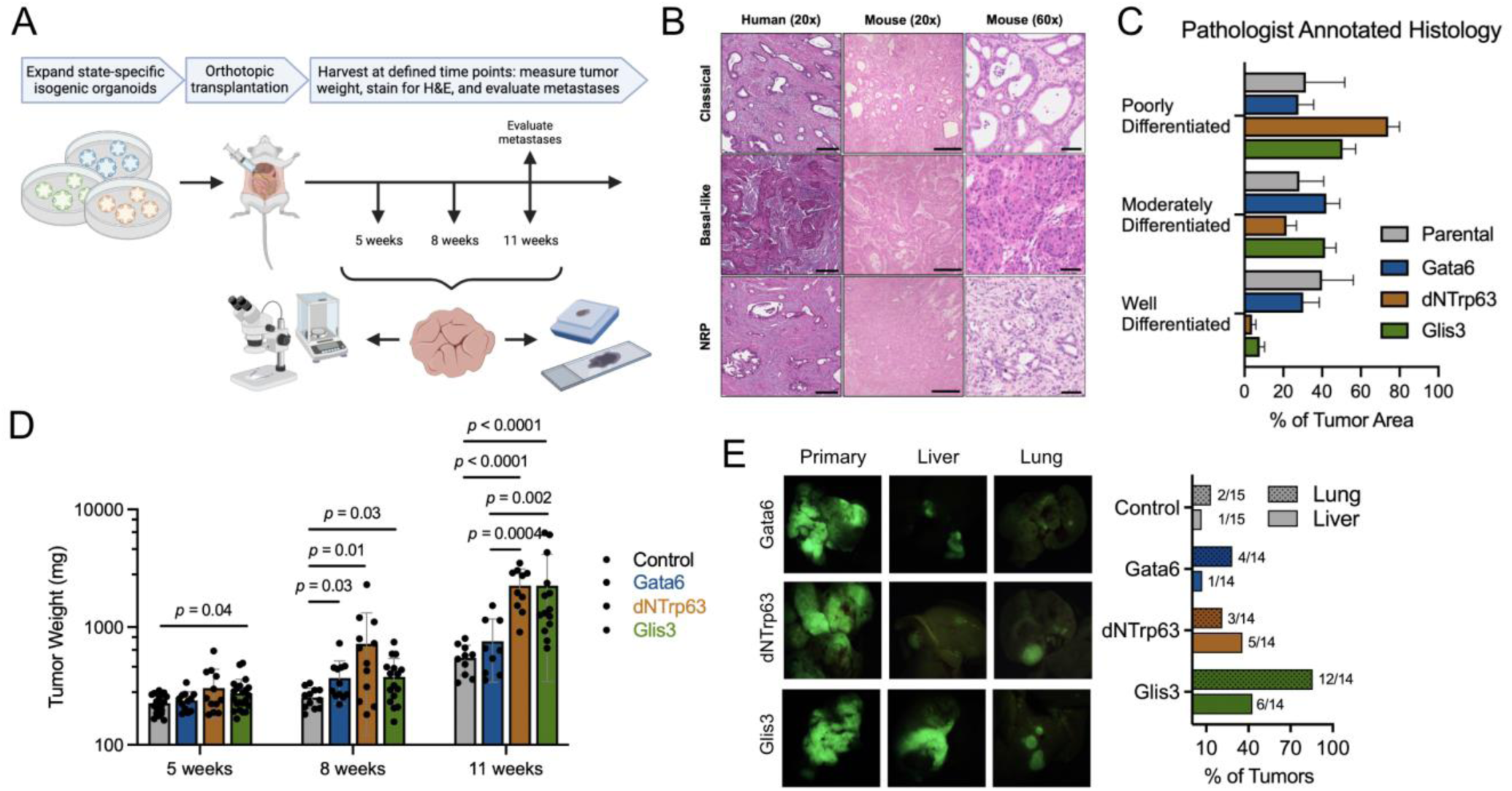
Isogenic models reveal aggressive and metastatic phenotypes *in vivo* and *in vitro*. (a). Schematic of orthotopic transplantation study of isogenic cell state models to evaluate *in vivo* growth kinetics. (b) H&E images from human and mouse tumors representative of neural-like progenitor (NRP), basal-like, and classical phenotypes. Scale bars for 20x objective images represent 200 µm and scale bars for 60x objective images represent 50 µm. (c) Histologic grading of tumor differentiation by board-certified pathologists (N.J.C., J.N., M.L.Z., M.M.-K.) in tumors of parental, Gata6, Glis3, or dNTrp63 background. (d) Tumor weights for mice transplanted with tumoroids at 5 weeks, 8 weeks, and 11 weeks. (e) Image panel of mNeonGreen+ tumors from orthotopic transplantation at the primary site (pancreas), liver metastases, and lung metastases (*left*) and bar plot showing quantification of metastatic frequency to the liver and lung from each isogenic model (*right*).

At 5 weeks, there was a significant growth advantage (*p* < 0.05; Mann–Whitney U test) of the *Glis3*-activated tumor relative to the control tumor (sgRNA targeting *Olfr102*, an unexpressed gene), but no significant differences among tumors derived from the other lines (**Fig. 5D**). At 8 weeks, we observed a significant growth advantage (*p* < 0.05; Mann–Whitney U test) of all cell state models (*Gata6*, *dNTrp63*, and *Glis3*) relative to the control (**Fig. 5D**). At 11 weeks, there was no longer a significant difference between the CLS and control tumors, but we continued to note significantly larger NRP (*p* < 0.0001; Mann–Whitney U test) and BSL (*p* < 0.0001; Mann– Whitney U test) tumors *vs.* control tumors (**Fig. 5D**).

In addition to differential primary tumor growth kinetics, we also observed pronounced differences in the metastatic proclivity of each malignant cell state. Substantiating observations made of CLS tumors in patients, *Gata6*-driven CLS tumors metastasized more frequently to the lungs (28.6%; 4/14 tumors) than the liver (7.1%; 1/14 tumors) (**Fig. 5E**). The *dNTrp63*-driven BSL model metastasized more frequently to the liver (35.7%; 5/14 tumors) than lungs (21.4%; 3/14 tumors) (**Fig. 5E**), which was also consistent with enrichment of the BSL state in human liver metastases^8^. Most notably, we observed the highest frequency of both liver and lung metastases in the *Glis3*-driven NRP model. Nearly half (42.8%; 6/14 tumors) of *Glis3* tumors metastasized to the liver, and most (85.7%; 12/14 tumors) metastasized to the lungs (**Fig. 5E**).

To further investigate gene expression patterns in the NRP primary tumor versus metastatic sites, we performed snRNA-seq on matched primary, liver, and lung tumors from the same *Glis3* mouse to control for confounding variables across different animals and extracted annotated malignant epithelial cells for comparison (**Fig. S9A**). Both lung and liver metastases had an abundance of differentially expressed genes relative to the primary tumor, in addition to sustained overexpression of *Glis3* (**Fig. S9B**). Genes enriched across both metastatic sites related to interferon and viral response signaling (*p* = 9.22e-8) (**Fig. S9C**), while genes enriched in the primary tumor related to integrin and focal adhesion signaling (*p* = 1.86e-6), TGF beta signaling (*p* = 1.32e-4), and HIF1a signaling (*p* = 4.06e-4) (**Fig. S9D**).

In mice, Glis3-driven organoids seeded a greater number of metastatic lesions compared to Gata6- and dNTrp63-driven organoids (**Fig. 5E**). However, in a separate cohort of PDAC patients with matched liver metastases and primary tumors^53^, we did not observe higher NRP expression in metastatic liver metastases compared to the primary tumor (**Fig. S9E**). To further study the relationship between NRP genes and metastasis, we queried the Cancer Metastasis Map^49^, containing metastatic proclivity data of over 450 human cancer cell lines. Here, we found NRP genes were expressed higher in strongly metastatic cell lines relative to non-metastatic cell lines, with *Glis3* featuring a particularly strong association with metastasis (**Fig. S9F**). Hence, we concluded that while Glis3 confers a pronounced metastatic phenotype and constitutive overactivation of Glis3 in a malignant genetic background can sustain NRP expression in metastatic sites, in human tumors, certain pancreatic tumor microenvironment features important for sustaining the NRP phenotype may be absent in distant metastatic sites, leading to decreased expression of the NRP state after the metastatic cascade has occurred.

## Discussion

Mounting scientific evidence suggests that diverse coordinated gene expression programs, or transcriptional cell states, may have a sizable impact on clinically relevant outcomes in cancer patients. Indeed, it has been established that classical and basal-like PDAC tumors exhibit differential prognoses and response rates to multi-agent chemotherapy^9^. More recently, we discovered a novel neural-like progenitor (NRP) malignant phenotype^7^ enriched after neoadjuvant cytotoxic treatment that has yet to be characterized in detail. In this study, we demonstrate that this residual post-treatment phenotype arises from treatment induced tissue-repair programs that result in overexpression of developmental transcription factors. One specific factor, GLIS3, enables residual cancer cells to rapidly repopulate local and distant sites, recapitulating the clinical pattern of increased metastatic disease at recurrence.

We show that NRP signatures are strongly enriched in regenerative post-injury mouse pancreata, both oncogene transformed and wild type. In humans, we discover that NRP is expressed by a stressed stem-like ductal progenitor population in physiologic pancreata and a DCLK1 expressing cancer cell population with poorly differentiated cellular morphology in malignant pancreatic tumors. Spatial colocalization analysis nominated inflammatory CAFs and IL-1β+ macrophage populations as producers of inflammatory factors such as IL-1β that upregulate NRP genes in cancer cells. Taken together, our results suggest that an inflammatory niche in the tumor microenvironment maintains NRP expression *in vivo*, which is absent with *in vitro* monoculture and may explain the difficulty in maintaining the NRP phenotype in culture.

Moreover, we found that cells utilizing the NRP program are marked by co-expression of transcription factors involved in pancreatic development, and we identified Glis3 as a transcription factor highly expressed in NRP cells and induced by IL-1β treatment. Isogenic organoids overexpressing Glis3 through CRISPR activation have markedly higher colony formation potential, increased invasive and migratory capacity, and metastatic proclivity in the orthotopic transplant setting relative to Gata6- and dNTrp63-driven classical and basal-like lines, respectively. Further evidence of Glis3 having a role in metastasis was gleaned from the observation that Glis3 expression was higher in human PDAC metastases relative to matched primary tumors.

Our data suggests several key questions for further study. In particular, it remains unclear whether the expression of neural genes by cancer cells mediates interactions with the nervous system, such as coordinating neurogenic tissue repair or recruitment of intratumoral nerves, or whether expression represents a broader ductal dedifferentiation program. For example, another study identified Tuft cell dedifferentiation dependent on *Myc* activity as a source of the NRP identity^54^. Stress and inflammation-mediated dedifferentiation has been reported extensively in the physiologic pancreas, with stressed ductal cells expressing neuroendocrine markers and progenitor features^55^. In some instances, this lineage plasticity in both physiologic and malignant ductal cells is dependent on the transcription factor STAT3^56,57^ downstream of IL-6 signaling, which we had previously reported to be enriched in the PDAC tumor microenvironment after CRT^32^. Several other studies have investigated the plasticity inducing effects of IL-1β and its role in epithelial regeneration. For example, AT2 cells in the lung respond acutely to IL-1β and Hif1α signaling pathways by increasing plasticity during injury^58,59^ and redifferentiate after suppression of inflammation and/or KRAS signaling^60^. Further study of residual disease in the post-treatment microenvironment may provide greater clarity behind the role of KRAS in maintaining the damage associated plasticity in PDAC, especially in light of recent findings supporting the efficacy of combining chemotherapy and KRAS inhibition^61^.

Our study has important limitations. Due to the scarcity of paired pre- and post-treatment biopsies, our study relies mostly on observations from unmatched patient samples. Additionally, while our experiments suggest that there exists a temporal component to expression of the NRP program, we are unable to directly confirm this in patient samples. We note that our TF driven isogenic organoid models still do not completely recapitulate the transcriptional properties of cell states in human PDAC tumors, with only a subset of each respective transcriptional signature being overexpressed along with the CRISPR activated driver. More faithful expression of the entire signature may necessitate specific microenvironmental factors. Furthermore, multifactorial perturbations may be required to fully recapitulate *in vivo* transcriptional phenotypes. While Glis3 seemingly drives several malignant phenotypes following cytotoxic therapy, cell states are heterogeneous and likely not captured by the functions of a single regulon.

Overall, this study deepens our understanding of residual disease in PDAC and provides functional characterization of the neural-like progenitor cell state, which is present in both primary tumors and metastatic disease and enriched following chemotherapy and radiotherapy. We discovered that injury- and inflammation-associated IL-1β signaling plays an important role in activating and maintaining the NRP phenotype in conjunction with rest and recovery after cytotoxic treatment. We identified the role of the developmental transcription factor *GLIS3* in driving the NRP phenotype and demonstrated in preclinical *in vitro* and *in vivo* models that *GLIS3*-driven NRP cancer cells are highly clonogenic, invasive, and metastatic. Taken together, NRP is an aggressive cancer cell phenotype that arises from treatment induced reprogramming, underscoring the clinical importance of identifying and targeting this malignant cell state in patients.

## Methods

### Isolating Organoids from Pancreatic Tissue

The mouse was genotyped to confirm Kras^LSL-G12D/+^;Trp53^FL/FL^;R26-dCas9-VPR-mNG. At around 6 weeks, the mouse was euthanized via isoflurane overdose and the pancreas was promptly removed onto a plastic dish. A razor blade was used to chop the pancreas into homogeneous tiny pieces, and then resuspended in 1 mL of digestion buffer consisting of 1x collagenase IV in PBS. The mixture is moved to an Eppendorf tube and placed into a rotating hybridization oven set to 37C for 20-30 minutes. After incubation, the tube was moved into a sterile tissue culture hood and the mixture was pipetted through a 70 µm filter placed over a 50 mL conical tube. An additional 40-50 mL of PBS was poured over the strainer. The tube was spun at 800G for 1 minute with acceleration at 9 and deceleration at 3. The supernatant was carefully aspirated and the pellet was resuspended in 1 mL PBS. The mixture was moved to a 15 mL conical tube and spun at 800G for 1 minute with acceleration and deceleration set to 9. The supernatant was carefully aspirated and the pellet was resuspended in 210 µL Matrigel (Corning, 356231). The solution was then pipetted into seven 30 µL domes into a 6-well plate (GenClone, 25-105). The domes were left to set in a 37C incubator for 20 minutes before 3 mL of culture media was added on top. The composition of the culture media was previously described in *Raghavan et al. 2022*^8^, consisting of 10 mM HEPES (Gibco), 500 nM A83-01 (Tocris), 100 ng/mL mNoggin (Peprotech), 10 nM hGastrin I (Sigma), 1x Primocin (Invivogen), 10 mM Nicotinamide (Sigma), WNT3A conditioned media 50% final, RSPONDIN-1 conditioned media 10% final, Advanced DMEM/F12 (Gibco), 100 U/mL penicillin/streptomycin (Gibco), 1x GlutaMAX (Gibco), 50 ng/mL mEGF (Peprotech), 100 ng/mL hFGF10 (Peprotech), 1.25 mM N-acetylcysteine (Sigma), and 1x B27 supplement (Gibco). We refer to this media composition as OPAC. The organoids were passaged a couple times before proceeding to spinfection with Ad-Cre.

Human organoids were derived in a similar manner, as previously detailed in *Raghavan et al. 2022.* In brief, PDAC tissue specimens were chopped and digested in the media described above with the addition of 10 µM Y27632 (Selleck), 1 mg/mL collagenase XI (Sigma Aldrich), and 10 ug/mL DNase (Stem Cell Technologies). After dissociation, the cells are seeded in Matrigel and cultured in media containing 10 μM Y27632 (Selleck) until the first media change. Human organoids may take longer to establish compared to murine organoids, often requiring multiple passages before they are robust enough for experimentation.

### Virus Production

CRISPRa and CRISPRko sgRNA sequences were designed using CRISPick by Broad Institute. We chose the SpyoCas9 (NGG) enzyme, and picked the top sequence produced per gene of interest. From there, we generated the reverse complement sequence, added “CACCG” to the beginning of the guide sequence, added “AAAC” to the beginning of the reverse sequence, and “C” to the end of the reverse complement sequence. The following primer pairs were ordered through IDT. We used the lentiGuide-Puro plasmid (Addgene, #52963, RRID:Addgene_52963) and followed the corresponding target guide sequence cloning protocol to produce our vector plasmids. The other components VSVG (Addgene, #8454) and PsPax2 (Addgene, #12260) were ordered and amplified through standard miniprep protocol. HEK293T cells (Takara Bio, 632180, RRID:CVCL_0063) were plated in 15 cm dishes to reach ∼70% confluency the following day. For each gene target, 2 mL of Opti-MEM (Gibco, 31985062) was mixed with 10 µg vector, 7.5 µg PsPAX2, 2.5 µg VSVG, and 60 µL TransIT-LT1 (Mirus, MIR 2306). This mixture was incubated at room temperature for 30 minutes, and then gently pipetted over the plate of HEK293T cells in 20 mL fresh DMEM + 10% FBS. After 72 hours of incubation at 37C, the supernatant is filtered through a 0.45 µM syringe filter and concentrated to a final volume of 1.6 mL using 10KDa Amicon Ultra-15 Centrifugal Unit (Millipore Sigma, UFC901024). The concentrated virus is aliquoted and stored at -80 until use.

### Spinfecting Organoids

Day 4-5 organoid domes were manually dissociated with a P1000 pipet and moved to a 15 mL conical tube filled with PBS. The tubes were spun at 800G for 1 minute and the pellet was resuspended in 2-3 mL of TrypLE Express (Gibco, 12604013). The tube was placed in a 37C incubator and manually agitated with a P1000 pipet every 10 minutes for a total of 30-40 minutes. Once the organoids were dissociated into single cells, PBS was added to fill the tube, spun to pellet, resuspended in culture media, and counted using a hemocytometer. 75k cells were loaded into each well of a 24 well plate (GenClone, 25-107). Appropriate amounts of the virus were added into each well and culture media was added until total volume reached 500 uL. We typically added 3 µL for Ad-Cre (U of Iowa-5, Ad5CMVCre) and 100 µL for our concentrated sgRNA lentivirus. The plate was spun at 600G for 1 hour at 23C, returned to the 37C incubator for 6 hours, and then replated in seven 30 µL Matrigel domes into a 6-well plate (GenClone, 25-105). Selections such as nutlin and puromycin were added to the media two days after spinfection.

### qPCR validation

RNA was isolated via the Invitrogen TRIzol Plus RNA Purification Kit (12183555) and quantified via NanoDrop. We loaded 1 µg of template RNA for cDNA construction to maintain consistency. cDNA is created and amplified via the Thermo Fisher SuperScript IV First-Strand Synthesis System (18091050). Primer sets were ordered through IDT under PrimeTime Predesigned qPCR Assays. We chose the top recommended assay with the highest algorithm score whenever possible, and we chose the “intercalating dyes, primers only” configuration. We ran qPCR using Sybr Fast 2x MM LC480 from Kapa Biosystems (KK4611) and the Bio-rad CFX384 Touch Real-Time PCR Detection System. Each sample had four technical replicates and Gapdh/GAPDH was used as the housekeeping gene. The relative fold difference was calculated using the delta-delta Ct method.

### Multiplex Immunofluorescence Imaging

Organoids were plated in a chambered slide (Ibidi, 80841) following dissociation at a sensity of 1,000 cells per 10 µl 70% Matrigel in OPAC and allowed to grow for five to seven days. Alternatively, organoids can be grown in dishes and smeared onto a slide. Two stock buffers were made: 10x PBS/glycine (38g NaCl, 9.38g Na2HPO4, 2.07g NaH2PO4, and 37.5g glycine in 500 mL DI Water, pH7.4, filtered) and 10x IF wash (38g NaCl, 9.38g Na2HPO4, 2.07g NaH2PO4, 2.5g NaN3, 5g BSA, 10 mL Triton X-100, and 2.5 mL Tween-20 in 500 mL DI Water, pH 7.4, filtered). The organoids were fixed in 2% PFA for 45 minutes at room temperature and then washed in 1x PBS/glycine for 10 minutes three times. After washing in 1x IF wash for 10 minutes three times, slides were blocked in primary block (1x IF wash + 10% goat serum) for two hours at room temperature. Primary antibody cocktail (diluted in primary block) was added for overnight incubation at 4 °C. After three washes, slides were incubated with Alexa Fluor– conjugated secondary antibodies (1:1000 in 1 × IF wash + 5 % goat serum) for 1 h at room temperature in the dark, and washed again 3 × 10 min. Nuclei were stained with DAPI (1 µg/mL, 5 min), rinsed twice in PBS, and mounted with ProLong Glass antifade mounting media prior to imaging.

### PDAC tumor production via orthotopic transplant

Day 4-5 organoid lines were dissociated to single cells and resuspended into 50% Matrigel 50% culture media at a concentration of 1k cells/uL. This solution was transferred into a 1 mL syringe using a 18G needle and placed into an empty 15 mL conical submerged in ice to prevent solidification. Our subject selection criteria were syngeneic female and male mice (P53 fl/fl; R26-LSL-dCas9VPR-mNG) at 8-12 weeks of age and healthy weight, were randomly allocated to each organoid group and prepped for surgery. The pancreas was exposed through a small ventral midline incision, and ∼100 µL (100k cells) was injected into the pancreas via a 30G needle. The wound was sutured close and the animal was sacrificed at the specified post-operation endpoint (5 weeks, 8 weeks, 11 weeks, moribund). During the harvest, the organs of interest (pancreas, spleen, lung, liver) were removed, weighed, and imaged for tumor growth in a single-blind manner. The tumor specimens were then prepped for downstream experiments. Parts of the tumor were flash frozen for single-nucleus RNA sequencing, whereas the remaining parts were fixed in formalin at 4C overnight followed by paraffin embedding, sectioning, and H&E. Animals that died prior to specified post-operation endpoints were excluded from all analyses. All procedures were approved by the Institutional Animal Care and Use Committee of Massachusetts General Hospital (protocol #: 2022N000162).

### 2D Cell Culture

Cell lines adapted for 2D culture were generated by culturing dissociated organoids in Advanced DMEM/F12 with 10% fetal bovine serum and 100 U/mL penicillin/streptomycin over the course of 2-3 weeks with serial passaging. Cell lines had distinct cellular morphology and were authenticated by RNAseq at derivation to share similar transcriptional state as their parental 3D counterparts. We utilized cells with passage number less than 30 for all experiments and did not observe shifts in morphology or behavior past this point. Mycoplasma contamination was performed every 2 months.

### Colony Formation Assays

In the 2D setting, cells were detached using TrypLE and seeded in 6 well plates at a density of 500 cells/well. After 7 days, cells were stained using crystal violet staining solution composed of 0.125 grams of crystal violet in 50 milliliters of 20% methanol. Colonies were imaged against a bright white background and quantified using the ‘Analyze Particles’ module in ImageJ (RRID:SCR_003070) to measure average colony diameter, covered plate area, and total colony count within each well. For 3D assays, organoids were dissociated into single cells and resuspended in 70% Matrigel 30% culture media at a concentration of 500 cells per 10 uL, which represented the volume of each seeded Matrigel dome. Seven technical replicates were seeded in each well of a 12 well plate (GenClone, 25-101). After 10 days, organoids were imaged on a light microscope at 10x magnification. All organoids were counted and measured manually using ImageJ for a total of 7 replicates per cell line.

### Invasion Assays

Radial invasion assays were performed in two formats, one in 100% Matrigel and one in compressed collagen I matrix. These assays differ by a 40 fold difference in stiffness (0.04 kPa vs 1.5 kPa) the latter of which with compressed collagen I is similar to tumor tissue. The collagen I invasion assay follows the protocol for centrifuged collagen hydrogels described previously^62^. Cells were seeded as 30 µL domes at a concentration of 25k/dome in the Matrigel assay and 75k cells/dome in the stiff collagen I assay. Invasion was measured at 4 and 14 days respectively via imaging. The invaded area was quantified using ImageJ. We additionally performed transwell Boyden invasion assays to assess invasion and migration capabilities in each of our cell lines. Cells were serum starved for 24 hours and then seeded at 50,000 cells per Matrigel coated insert (Corning 354480) with 500 µl serum containing media below the insert and 500 µl serum free media containing cells seeded on top of the insert. Cells were incubated and allowed to invade for 48 hours and stained according to Diff-Quik staining kit instructions. Invasiveness was quantified by taking brightfield images of 5 representative fields of view and quantifying stained areas within the field as well as using the ‘Analyze Particles’ module in ImageJ to count cell bodies.

### 10x Single-nucleus RNA Sequencing

We followed the protocol detailed in *Hwang et al, 2022* with minor adaptations. In brief, a 2x stock of STc buffer with the final concentrations of 40 mM Tricine (VWR, E170-100G), 42 mM MgCl2 (Sigma-Aldrich, M1028), 292 mM NaCl (Thermo Fisher Scientific, AM9759), and 2 mM CaCl2 (VWR, 97062-820) was prepared in nuclease-free water. To prepare 3mL of 1x working STc buffer per specimen, we diluted the 2x STc buffer 1:1 in nuclease-free water. To prepare 2 mL of NSTcPA buffer per specimen, we combined 1 mL 2x STc buffer, 10 µl 2% bovine serum albumin (New England Biolabs, B9000S), 1 µl 1 M spermidine (Sigma-Aldrich, S2626-1G), 40 µl of 10% Nonidet P40 Substitute (Thermo Fisher Scientific, AAJ19628AP), 0.3 µl of 1 M spermine (Sigma-Aldrich, S3256-1G), 20 µL Protector RNase Inhibitor (Sigma, 3335402001), and 928.7µl nuclease free water. Additionally, we created a nuclei resuspension buffer consisting of PBS with 1:10 MACS BSA Stock Solution (Miltenyi Biotec, 130-091-376) and 1:40 Protector RNase Inhibitor. Utilizing tools cleaned with RNase Away (ThermoFisher, 7000TS1) followed by 70% Ethanol, the specimen was minced in 1 mL of NSTcPA buffer in a 6 well dish on ice. After approximately 8 minutes, the homogenized specimen was passed through a 40 µM Falcon cell filter (Thermo Fisher Scientific, 08-771-1) over a 50-ml conical tube. The well was rinsed with 1 mL of NSTcPA buffer, and 3 mL of 1x STc buffer was added. After spinning, the pellet was resuspended in the nuclei resuspension buffer at a volume to bring the final concentration to 300-2,000 nuclei per microliter. From this point onwards, the Chromium Next GEM Single Cell 3’ Reagent Kits v3.1 user guide was closely followed with an initial nuclei loading density of 10,000.

### Scoring of Public Datasets

We scored publicly available RNAseq and scRNAseq datasets from PDAC patients either untreated or receiving neoadjuvant chemotherapy (Human Tumor Atlas Network (HTAN) dbGaP Study Accession: phs002371.v1.p1^16^, GSE205013^17^, GSE156405^18^, GSE202051^7^, supplementary tables^19^), human metastatic PDAC specimens (dbGaP: phs001652.v1.p1^20^), human cell lines (https://depmap.org/portal/data_page/?tab=overview), mouse pancreatic spheroids derived from pancreas either pre-exposed or not exposed to inflammation (GSE180211^24^), mice treated with caerulein and analysed at acute timepoints (NCBI BioProject ID PRJNA978570^25^), KRAS^G12D^ background mice treated or untreated with caereulein (GSE207943^26^), the pancreatic progenitor cell niche (GSE131886^27^), laser capture microdissected morphology guided sequencing (GSE208732^28^), and spatial transcriptomics on patient PDAC specimens (Mendeley data: https://doi.org/10.17632/kx6b69n3cb.1). For these datasets raw count matrices or counts in supplementary tables were downloaded and analysed as described in the text. Using the TCGA web portal, we downloaded transcriptomic and clinical data from the pancreatic adenocarcinoma (PAAD) cohort for pancreatic cancer. Signature scores were calculated using the scanpy.tl.score_genes module in Scanpy for scRNAseq datasets and using ssGSEA for TPM transformed RNA sequencing datasets. Within each dataset, NRP score was Z-normalized.

### RNA sequencing analysis

For the chronic ligand treatment experiments, organoids were seeded at 1,000 cells per 30 µL dome in 6 well plates as previously described. Cells were continuously treated in refreshed media every two days and passaged over the course of 3 weeks. We used mouse recombinant IL-1β (Thermo Fisher 211-11B-10UG) at a concentration of 10ng/mL, mouse recombinant hyper IL-6 (R&D Systems 8954-SR-025/CF) at a concentration of 20 ng/mL, and 5-FU (Sigma F6627-1G) dissolved in DMSO at a concentration of 3 µM. For 5-FU monotherapy experiments, we treated with 3 µM 5-FU for 7 days, refreshing media every two days. RNA was collected from organoids using Trizol (Invitrogen 15596026) according to manufacturer protocol and quantified with NanoDrop. RNA samples were sequenced with Novogene with Illumina PE150 reads (6 G raw data per sample). Raw FastQ files were processed and converted into counts files using Salmon and using hg38 as reference genome. Differential expression analyses were performed with DESeq2 (RRID:SCR_000154) in R v4.3.3.

### Quantification and statistical analyses

Statistical analyses were performed with GraphPad Prism v10.3.1 and R v4.3.3 in RStudio. Results are reported as mean ± s.d. or mean ± s.e.m and boxes in boxplots were plotted using the Tukey method. Significance was defined as P < 0.05. We supplemented traditional rank based GSEA analysis using predefined gene sets with an overrepresentation analysis method implemented by gProfiler (https://biit.cs.ut.ee/gprofiler/gost). Significantly enriched genes in a condition above a Benjamini Hochberg adjusted p-value threshold of 0.01 and a log2fc difference in means threshold of 0.25 were utilized to identify significantly enriched gene sets. For all gProfiler analysis, we capped the maximum gene set size to 500 genes. In general, sample size was based on estimations by power analysis with a level of significance of 0.05 and a power of 0.9.

## Acknowledgements

We thank the members of the Hwang and Aguirre lab for their helpful discussions and Nicole Lester, Dianne Moschella, Tina Balducci, Misha Pivovarov, Serena Sullivan, Sharon McSorley, Matthew Mues, Anthony Zietman, Daphne Haas-Kogan, Theodore Hong, Jennifer Wo, and Ralph Weissleder for scientific and administrative support. Additionally, we thank the following organizations for grant support: National Science Foundation Graduate Research Fellowship (DG), National Cancer Institute K08CA270417 (WLH), Burroughs Wellcome Fund Career Award for Medical Scientists (WLH), Pancreatic Cancer Action Network Career Development Award (WLH).

---

**Supplementary Figure 1:**
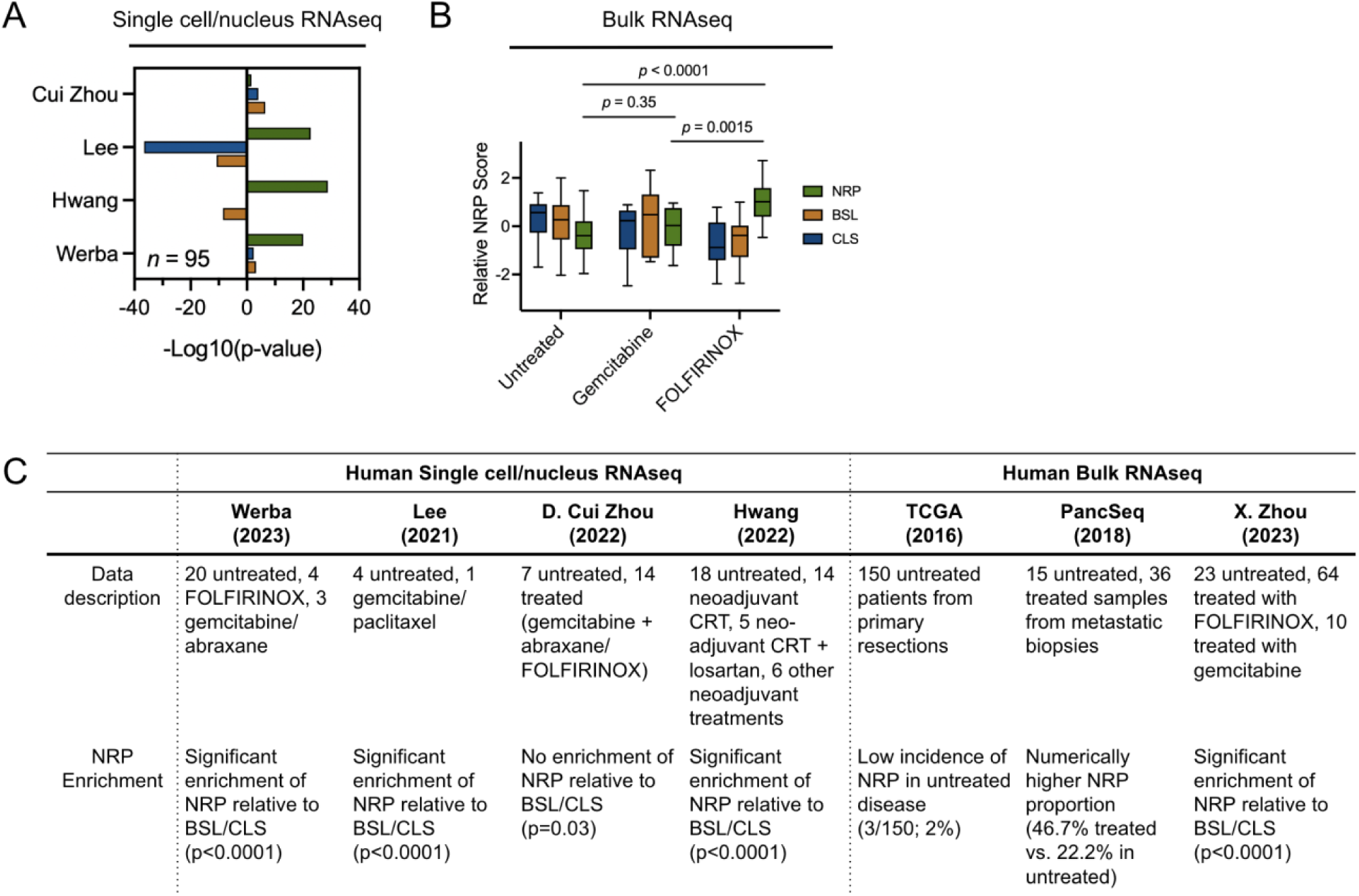
NRP scores in other PDAC datasets. (a) Grouped barplot depicting enrichment of NRP relative to classical and basal signatures in other independent scRNAseq cohorts^7,16–18^. (b) Boxplot of NRP score in bulk RNAseq samples of untreated specimens or those receiving neoadjuvant chemotherapy from the X. Zhou dataset. (c) Summary table of scored human datasets and their NRP enrichment in treated samples.

**Supplementary Figure 2:**
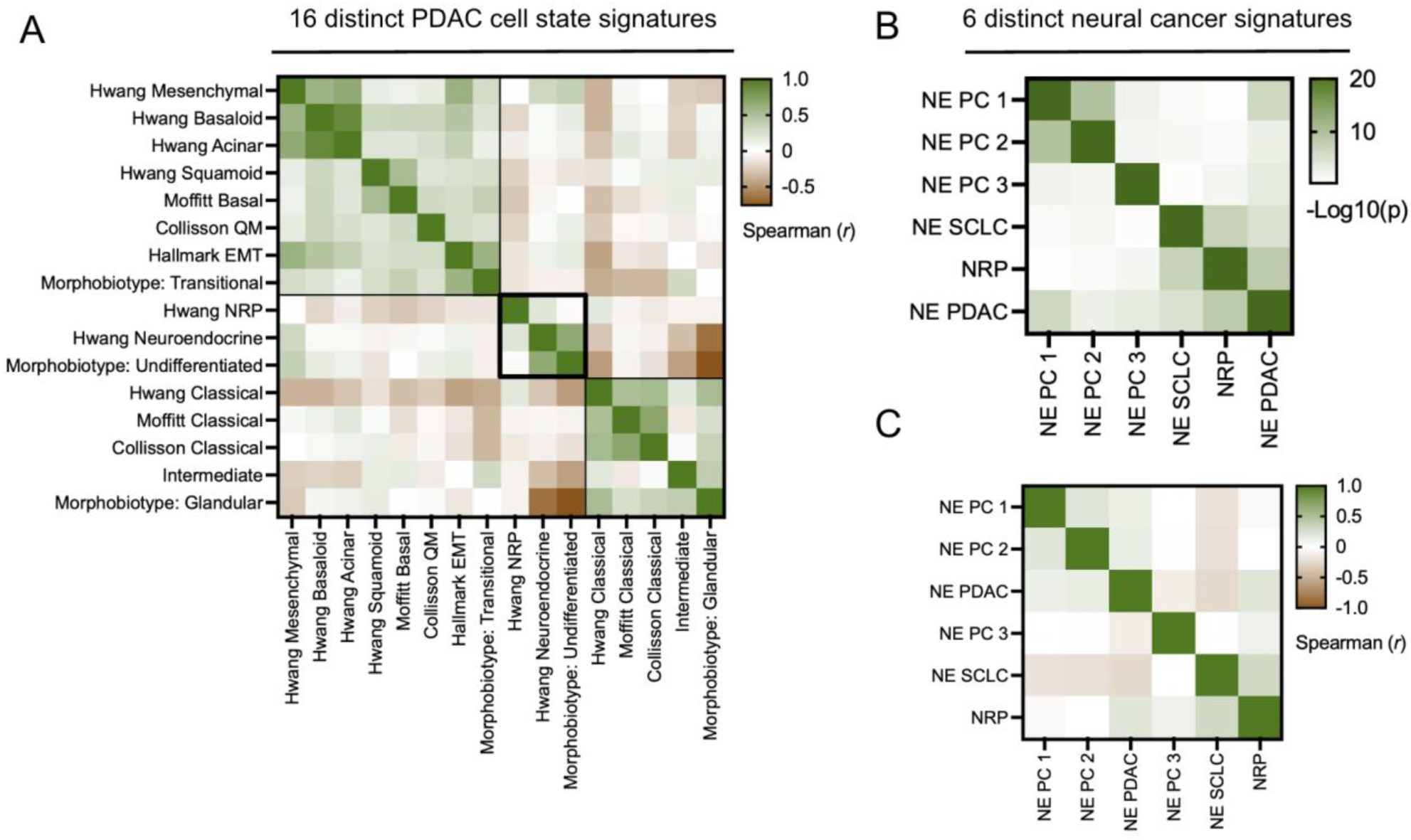
Comparison of neural-like progenitor state in PDAC to other transcriptional signatures in PDAC and neural-related signatures in other malignancies. (a) Clustered heatmap of Spearman correlation coefficient values between cell state signature scores across all annotated cancer cells in our snRNAseq data. Annotated groupings reflect similarity of various classical cell state signatures, basal-like cell state signatures, and neural or undifferentiated signatures. (b) Heatmap of negative log transformed p-values calculated via Fisher’s exact test between pairs of neural or neuroendocrine state signatures (NE PC 1 and NE PC 2 are two neuroendocrine prostate cancer signatures from the same study^63^, NE PC 3 derives from a separate neuroendocrine prostate cancer profiling study^61^, NE SCLC is a neuroendocrine small cell lung cancer signature^7^, and NE PDAC is a neuroendocrine PDAC signature^7^). Greater green intensity denotes increasing significance of gene set overlap. (c) Heatmap of Spearman correlation coefficients between neural or neuroendocrine state signatures across all annotated cancer cells in our snRNAseq data.

**Supplementary Figure 3:**
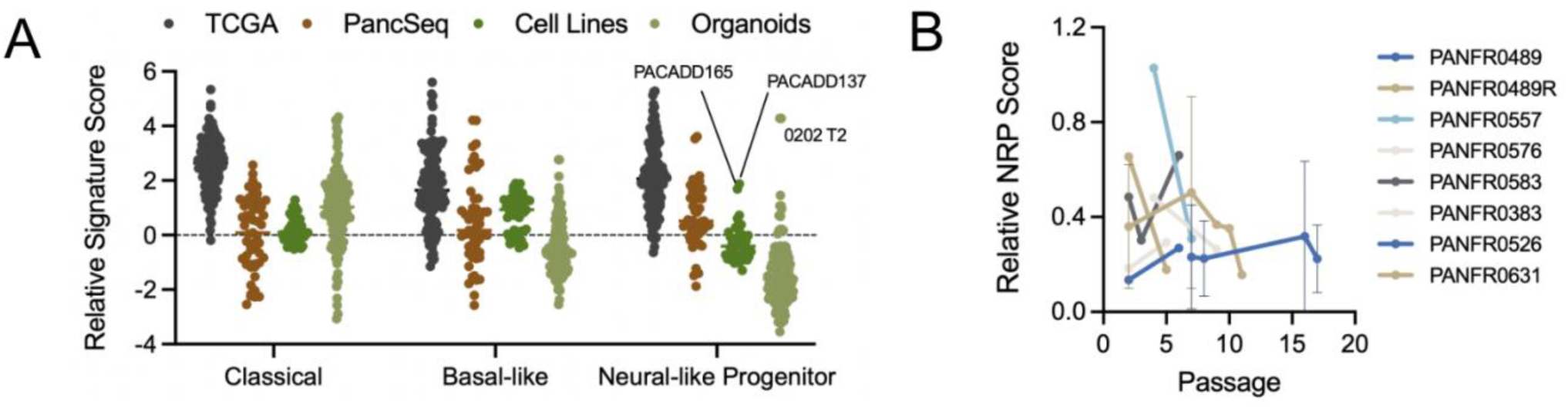
Poor representation of the neural-like progenitor cell state in conventional patient derived *in vitro* models. (a) Schematic and quantification of NRP score in patient tissues, patient derived organoids and 2D cell lines. (b) Quantification of NRP score in patient-derived PDAC organoid lines at serial time points during passage.

**Supplementary Figure 4:**
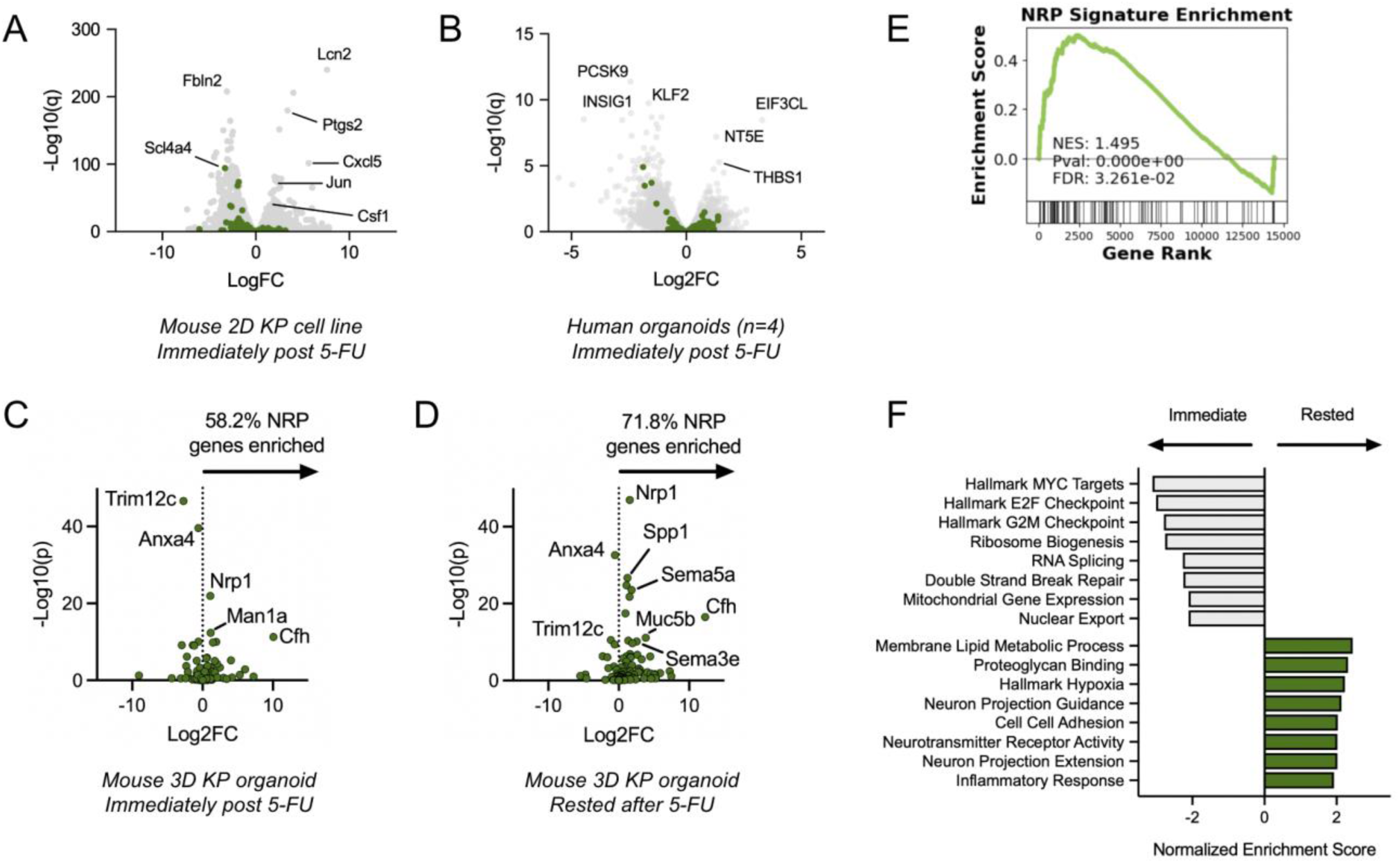
Chemotherapy induced genes in PDAC model systems. (a) Volcano plot showing differentially expressed genes in a mouse KP cell line after 48 hours treatment with 5-FU. (b) Volcano plot showing differentially expressed genes across 4 human organoid lines after 72 hours treatment with 5-FU. (c) Volcano plot showing differentially expressed genes in a 3D organoid culture system with mouse KP organoids immediately after 48 hours of chemotherapy relative to untreated cells. Volcano plot showing differentially expressed genes in a 3D organoid culture system with mouse KP organoids after 48 hours of 5-FU treatment, followed by 72 hours of resting. (e) GSEA plot of NRP gene set enrichment in rested cells relative to cells immediately after 48 hours of chemotherapy. (f) Grouped barplot of gene sets enriched in cells immediately after chemotherapy or after resting for 72 hours.

**Supplementary Figure 5:**
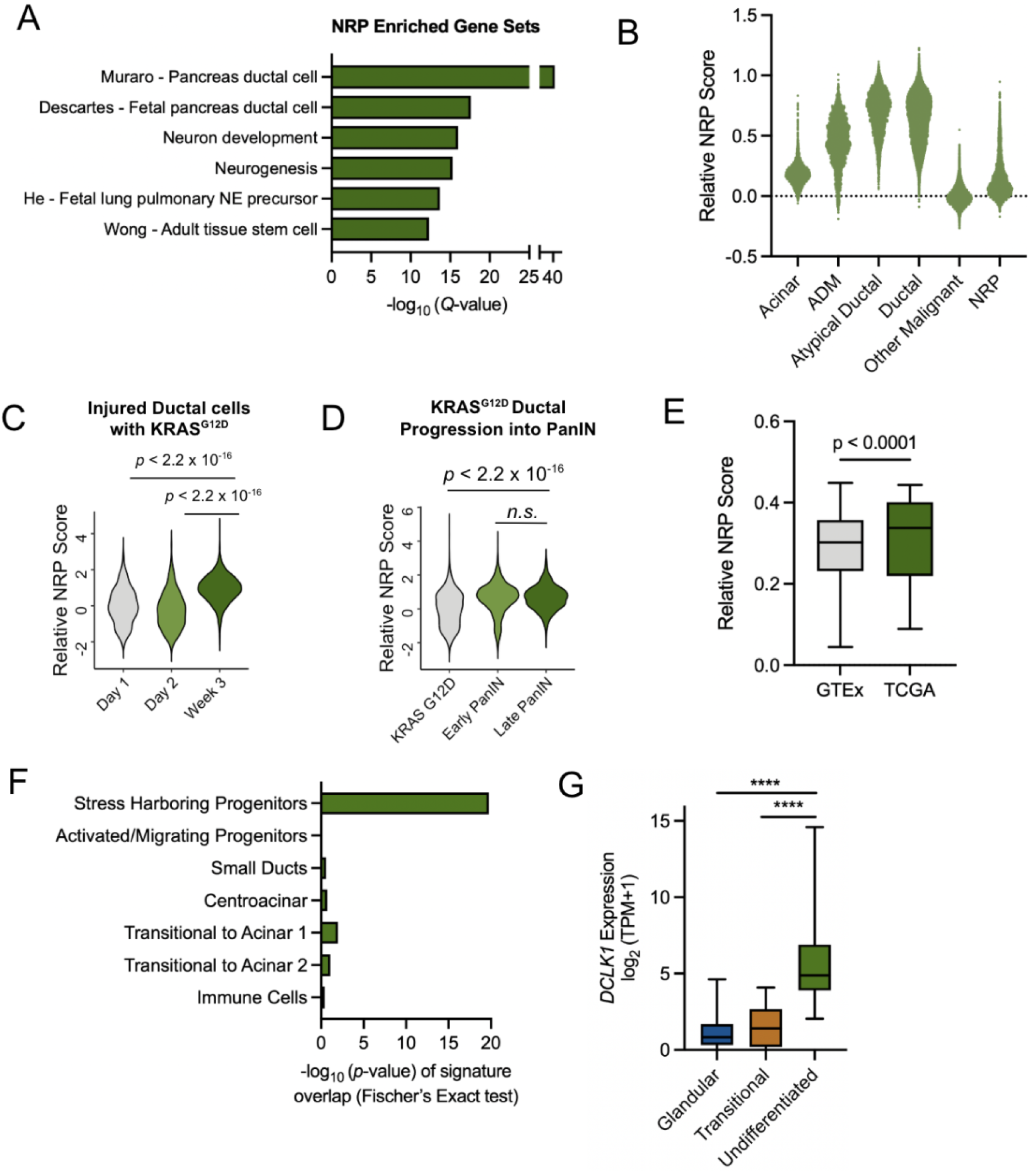
NRP scoring in pancreas cancer datasets. (a) Barplot of GSEA terms enriched in 200 gene NRP signature. (b) Jitterplot showing NRP score in annotated epithelial cell types from our previously published snRNAseq cohort^7^. (c) Violin plot of NRP score in Kras^G12D^ mutant ductal cells from cerulein-treated mice, assayed via scRNAseq. (d) NRP score through progression of Kras^G12D^ ductal cells to PanIN. (e) Boxplot showing NRP score in GTEx pancreas samples versus annotated normal pancreas samples from the TCGA. (f) Barplot depicting enrichment of NRP genes in pancreatic progenitor cell subsets from healthy human donors^27^ via Fisher’s Exact test. (g) *DCLK1* expression in microdissected undifferentiated cells by histology in PDAC patient samples^28^ which are enriched for neural-like genes.

**Supplementary Figure 6:**
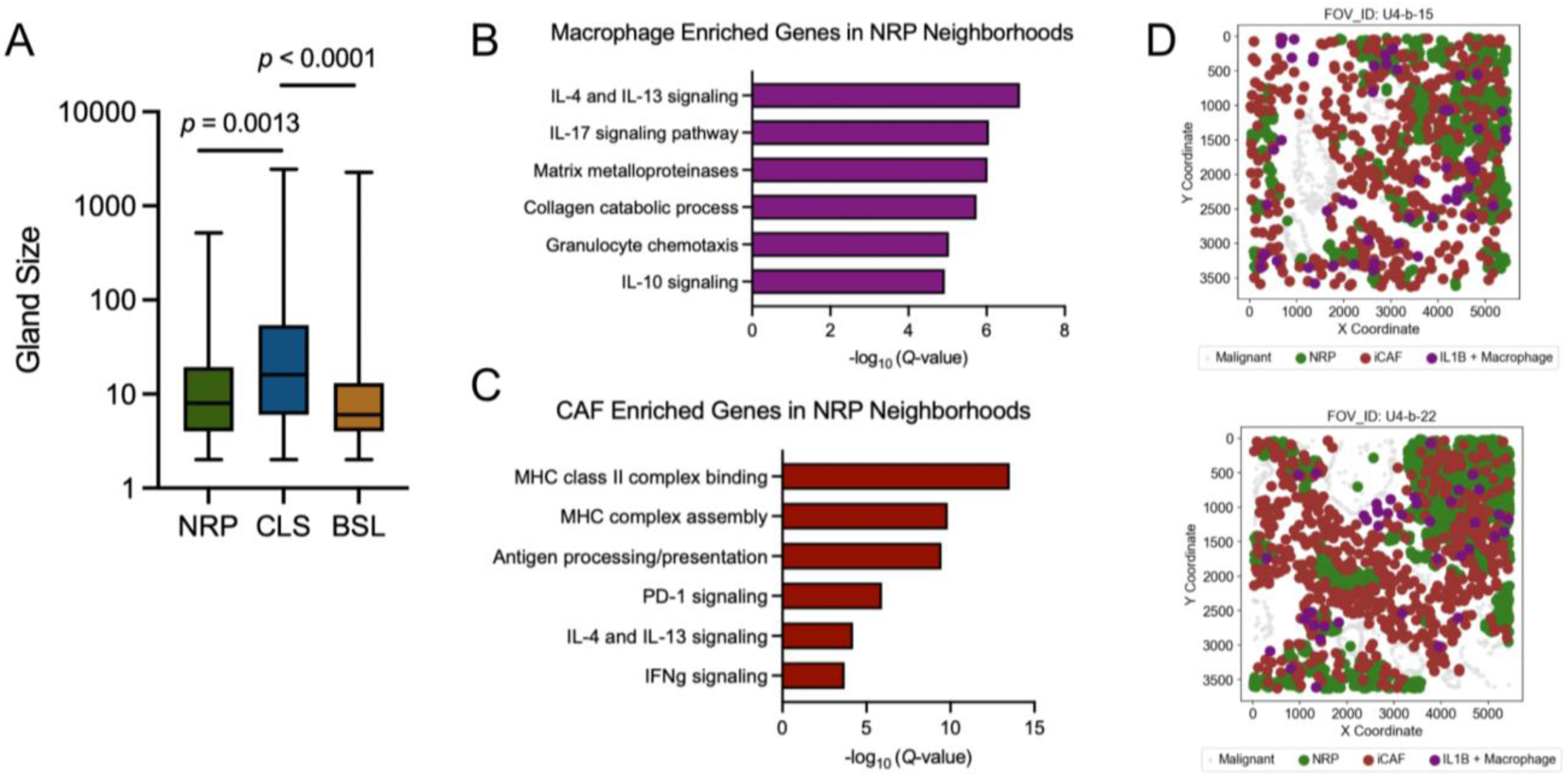
Neural-like progenitor expression programs are observed in pancreatitis and injury repair and enforced by the local microenvironment. (a) Quantification of gland sizes in glands dominated by BSL, CLS, or NRP cells. (b) Barplot showing enriched gene sets in CAFs around NRP cells relative to other CAFs. (c) Barplot showing enriched gene sets in macrophages around NRP cells relative to other macrophages (Methods). (e) Representative fields of view showing colocalization of NRP cells, iCAFs, and IL1β macrophages.

**Supplementary Figure 7:**
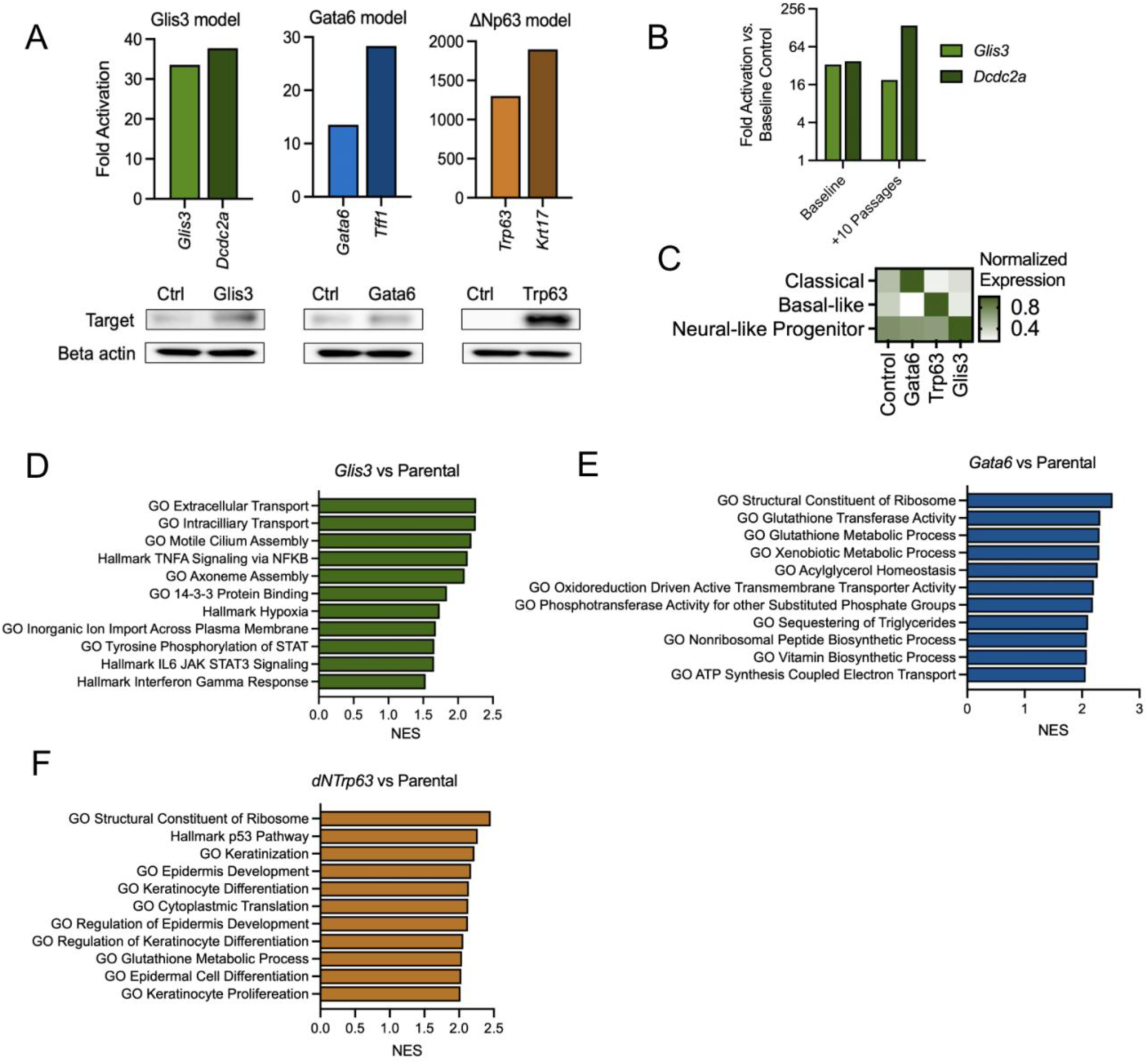
Generation and molecular characterization of isogenic models. (a) Validation of isogenic overexpression by qPCR and Western blot in organoid lines. (b) qPCR of select NRP genes after serial passage of isogenic organoids. (c) Heatmap depicting normalized relative signature score in isogenic organoids. (d) Barplot depicting overexpressed gene sets in Glis3, (e) Gata6, and (f) dNTrp63 isogenic overexpression lines.

**Supplementary Figure 8:**
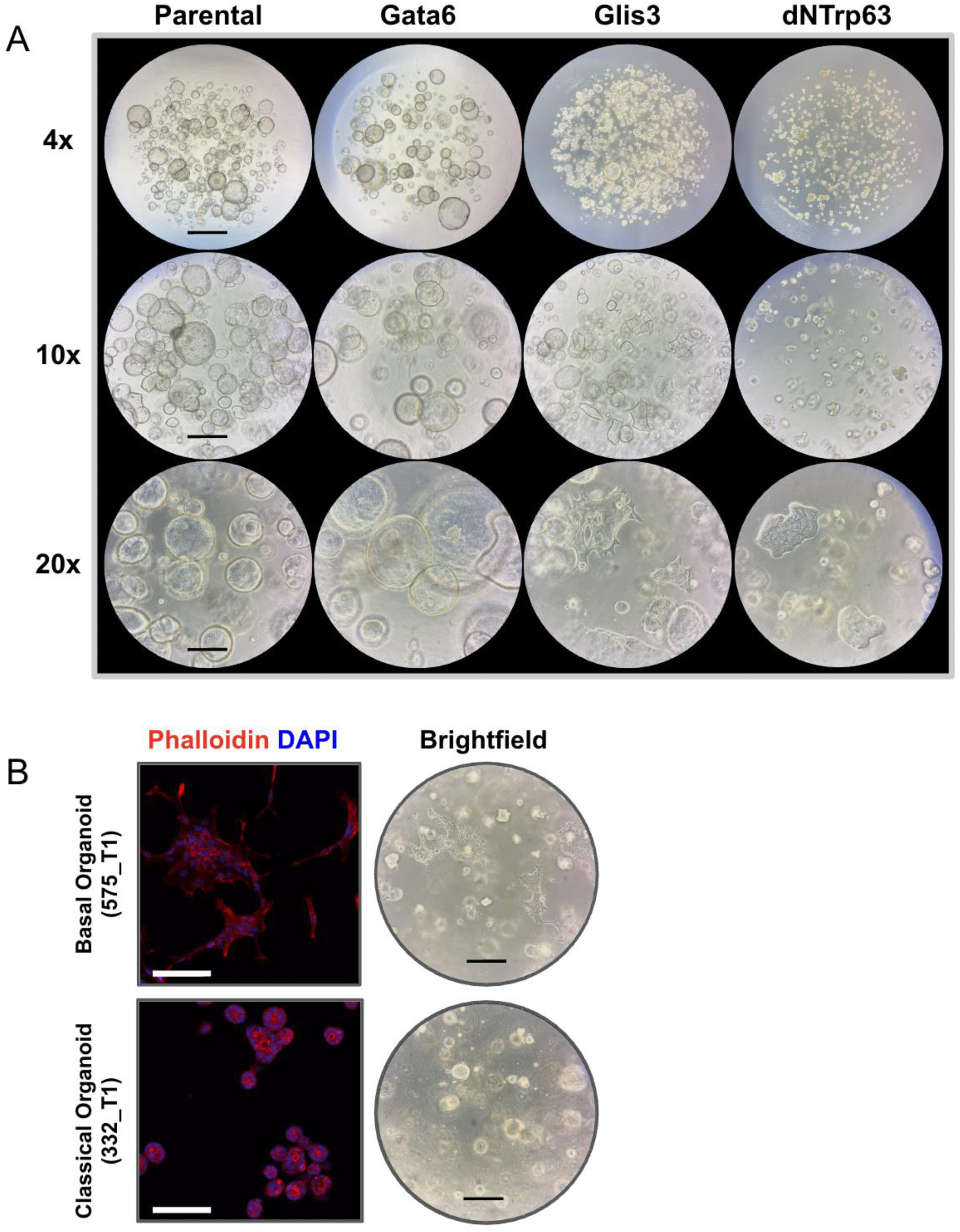
Brightfield imaging of isogenic organoids. (a) Phase imaging of isogenic organoids at magnifications of 4x, 10x, and 20x. Scale bars represent 1,000 µm, 400 µm and 200 µm respectively and the same scale is used across the magnification level. (b) Immunofluorescence and 10x brightfield imaging of human organoids classified as basal-like or classical by transcriptional signatures. Scale bars represent 100 µm at a 20x magnification.

**Supplementary Figure 9:**
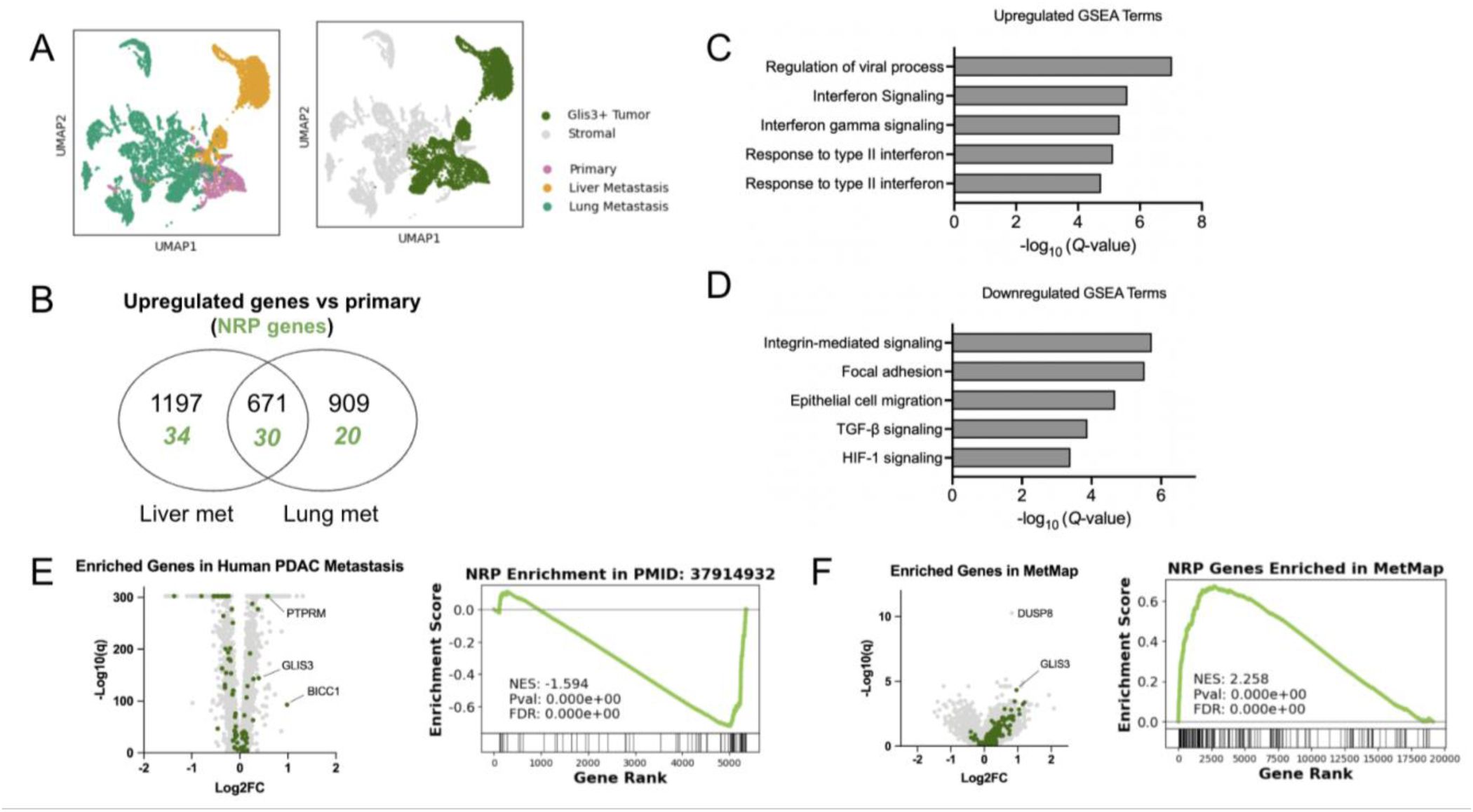
Neural-like progenitor signatures promote metastasis but expression is lost in metastatic tumor microenvironments. (a) UMAP projection plot of isogenic Glis3 matched primary and metastasis scRNAseq data colored by site (left) and cellular identity (right). (b) Venn diagram showing enriched genes in metastases relative to the primary tumor. (c,d) Barplot of GSEA terms upregulated (c) and downregulated (d) in metastases relative to the primary tumor. (e) Volcano plot highlighting NRP genes (green) enriched in human liver metastases in a cohort of matched samples profiled with scRNAseq^53^. (f) Volcano plot highlighting NRP genes (green) enriched in metastatic human cell lines as profiled by the MetMap project.

